# The Human Neural Cell Atlas of Zika Infection in developing human brain tissue: viral pathogenesis, innate immunity, and lineage reprogramming

**DOI:** 10.1101/2024.09.27.615512

**Authors:** Caleb Stokes, Leanne S. Whitmore, Dante Moreno, Karan Malhotra, Jennifer Tisoncik-Go, Emily Tran, Nick Wren, Ian Glass, Birth Defects Research Laboratory (BDRL), Jessica E. Young, Michael Gale

## Abstract

Zika virus (ZIKV) infection during pregnancy can lead to fetal brain infection and developmental anomalies collectively known as congenital Zika syndrome (CZS). To define the molecular features underlying CZS in a relevant human cell model, we evaluated ZIKV infection and neurodevelopment in primary fetal brain explants and induced pluripotent stem cell-derived mixed neural cultures at single cell resolution. We identified astrocytes as key innate immune sentinel cells detecting ZIKV and producing IFN-β. In contrast, neural progenitor cells displayed impaired innate immunity and supported high levels of viral replication. ZIKV infection of neurons suppressed differentiation and synaptic signaling networks and programmed a molecular switch from neurogenesis to astrogliogenesis. We identified a universal ZIKV-driven cellular stress response linked to intrinsic apoptosis and regulated by IFN-β. These findings reveal how innate immune signaling intersects with ZIKV-driven perturbations in cellular function to influence CZS outcomes including neuron developmental dysfunction and apoptotic cell death.

## Introduction

Zika virus (ZIKV) is an emerging mosquito-borne flavivirus that can cause fetal developmental anomalies upon infection during pregnancy. The features and outcomes of fetal ZIKV infection are collectively called congenital ZIKV syndrome (CZS) and range from developmental delay to severe microcephaly, but underlying mechanisms remain poorly understood^1^. As the endemic range of ZIKV is expanding with climate change^2,3^ and there are currently no approved treatments or vaccines to prevent infection, achieving a mechanistic understanding of ZIKV neuropathology is critical to improving outcomes in CZS.

In efforts to understand why ZIKV affects brain development so profoundly, many studies have focused on examination of neural stem cells (NSCs). Although NSCs are highly susceptible to ZIKV, the properties of NSCs that make them permissive to infection are not defined^4–8^. In postnatal brain infections, innate immunity plays a critical role in controlling viral spread wherein cells expressing pattern recognition receptors (PRRs) detect viral infection and produce cytokines, particularly interferon-beta (IFN-β), which induces expression of hundreds of antiviral genes. In contrast, embryonic stem cells neither produce nor respond to IFN-β^9–11^, indicating that the timing and function of innate immune maturation in the fetal brain are under discrete control. Studies of ZIKV infection in vitro have employed a wide range of cell types to model ZIKV infection of NSCs, yielding conflicting findings regarding innate immune activity^7,12–15^. Furthermore, recent work has shown that ZIKV has tropism across diverse neural cell types, including astrocytes^16–18^, neurons^19,20^, and oligodendrocytes^21,22^, in addition to NSCs. This broad neural cell tropism of ZIKV suggests viral disruption of phenotypes across multiple cell types may underlie CZS. However, no studies have directly compared the consequences of ZIKV across human neural cell types, nor is it known how ZIKV alters intercellular communication in developing fetal brain.

To define the molecular features of ZIKV infection and pathology across neural cell types, we performed single cell RNA sequencing (scRNAseq) of ZIKV infection in human primary fetal brain explants and in human induced pluripotent stem cell (hIPSC)-derived neural cells containing a diversity of neural cell types ^23,24^. We found that astrocytes expressed high levels of PRRs, and undergo robust innate immune activation and innate immune effector gene induction during infection, and are a major source of IFN-β. In contrast, NSCs failed entirely to undergo innate immune activation in response to ZIKV infection, revealing major differences in the neural cell response to ZIKV. Simultaneous virus-host bioinformatics analysis using scPathoQuant, which links host gene expression changes to ZIKV RNA levels in individual cells^25,26^, allowed us to distinguish innate immune and virus-driven neuropathology at single cell resolution. This analysis revealed a ZIKV-induced transcriptional response induced universally across infected neural cell types and mechanistically linked to virus-induced apoptotic signaling. Among all cell types, mature neurons had the highest number of ZIKV-related transcriptional changes, including suppression of genes controlling synaptic function and axon outgrowth. We also defined a virus-induced switch from neuron-specific to glial-specific differentiation programs, identifying a molecular basis for gliosis observed in CZS^27–29^.

Our data constitute the Human Neural Cell Atlas of Zika Infection at single cell resolution encompassing neural cell response to ZIKV infection and IFN-β. The atlas defines programs of ZIKV-induced innate immune activation and response, and neural cell-specific pathology in two tissue models of developing human fetal brain. The Human Neural Cell Atlas of Zika Infection can now serve as a resource to inform intervention strategies based on identified signaling pathways for preventing or repairing neurologic injury caused by congenital ZIKV infection.

## Results

### HUMAN FETAL BRAIN PRIMARY TISSUE, CELLS, AND ZIKV INFECTION

We isolated cortical tissue from human fetal forebrain explants at gestational ages ranging from 13 to 17 weeks post-conception (Fig. S1, Supplemental Table 1). This explant tissue spanned the full thickness of developing cortex from the NSC niche in the subventricular zone to the cortical plate (Fig. S1A-B). In two-dimensional cultures of dissociated primary fetal cortical tissue, immunocytochemistry (ICC) demonstrated diverse cell populations including NSCs, neurons, astrocytes, and oligodendrocyte lineage cells (Fig. S1C-E). We infected these mixed cultures with an Asian-lineage strain of ZIKV isolated from a patient Brazil in 2015 (ZIKV/Br), the height of the microcephaly epidemic, or a pre-epidemic Asian-lineage strain from Cambodia (ZIKV/Cam). While all cell types supported viral infection, NSCs and oligodendrocytes had the highest rate of ZIKV-positive cells by ICC (Fig. S1F).

### INTERFERON-BETA DOMINATES THE HUMAN FETAL BRAIN RESPONSE TO ZIKA INFECTION

We performed single-cell RNA sequencing (scRNAseq) of mixed neural cell cultures, using Garnett classifier to define cell type identities (Fig. S2, Fig. 1A-C). We analyzed cell-specific transcriptional changes after 48 hours of ZIKV infection, a time point corresponding to peak viral load (Fig. S1G). For controls, we treated cultures with viral growth medium alone (MOCK) or exogenous interferon-beta (IFN-β, 100 IU/mL for 6 hours). We distinguished NSCs, neurons (both excitatory and inhibitory), astrocytes, and oligodendrocyte lineage cells. We also identified microglia, but in numbers too small for downstream analysis. A small percentage (<1%) of cells contained reads aligning exclusively to the ZIKV genome, demonstrating ZIKV infection (Fig 1D, Fig S3)^25^. The relative proportions of ZIKV-positive cells identified in this manner were similar to those identified by ICC, indicating that ZIKV infects cells in primary fetal brain explants at low frequency, with viral RNA most abundant in NSCs, astrocytes, and oligodendrocytes (Fig. 1C, Fig. S3).

**Figure 1.**
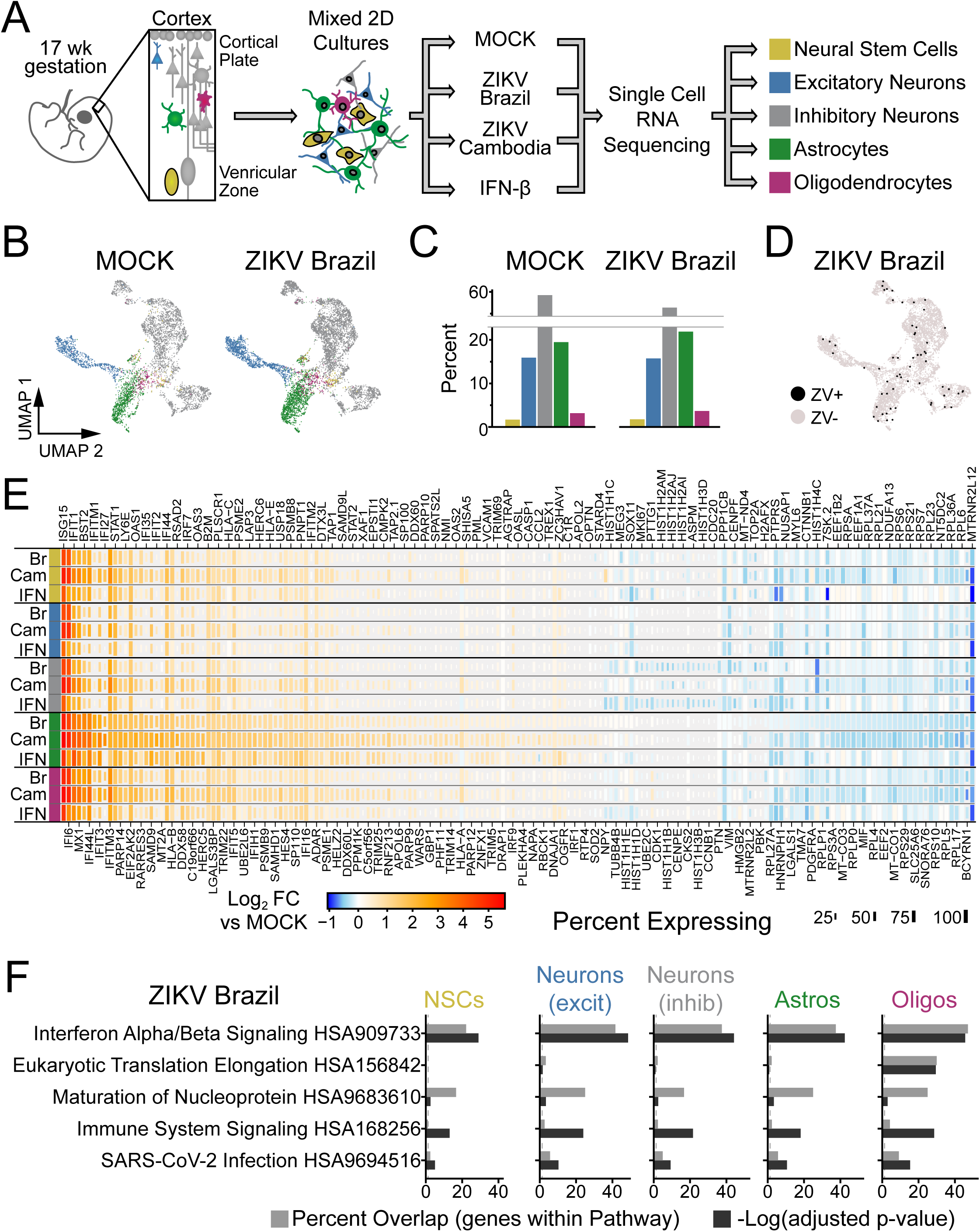
ZIKV infection of human primary fetal brain explants leads to extensive changes in gene expression driven by interferon-beta. (A) Experimental setup. Dissociated human primary fetal brain explant tissue (donor #1, 17 week male) was treated by infection with Zika virus (MOI=10) for 48 hours, mock infection, or IFN-β (100 μM for 6 hours). (B-C) Uniform Manifold Approximation and Projection (UMAP) plots (B) and relative percentage (C) of the predominant cell types identified in primary human fetal tissue. See Fig. S2 for additional conditions. (D) UMAP highlighting cells containing ZIKV reads (black) from ZIKV-Brazil condition. (E) Heat map (rows=treatment conditions, grouped by cell type) representing “pseudo-bulk” log_2_ fold change in gene expression relative to MOCK for the top 177 genes across cell types and conditions (|Log_2_FC|≥1.2, adjusted p-value <0.05 in at least one condition). (F) Overrepresentation analysis (ORA) identified type I interferon (α/β) signaling as the top pathway engaged by ZIKV-Brazil infection across cell types.

ZIKV infection caused extensive changes in gene expression that were highly similar across cell types (Fig. 1E, S4A, Supplemental Table 2). The pre-epidemic Cambodian ZIKV strain led to larger magnitude changes in differentially expressed (DE) genes despite fewer ZIKV-positive cells, consistent with previous work from our laboratory indicating Asian-lineage isolates induce a stronger innate immune response than American-lineage isolates associated with microcephaly^30^. Over-representation analysis (ORA) of these transcriptional profiles identified type I interferon (IFN-α/β) signaling as the most highly enriched pathway engaged during infection by both ZIKV isolates (Fig. 1F, Fig. S4B, Supplemental Table 3). Consistent with this outcome, gene expression changes due to ZIKV were highly correlated with those induced by exogenous IFN-β (Fig. S4C). These findings indicate that ZIKV-related transcriptional programs in human primary fetal brain explants are dominated by interferon-stimulated genes (ISGs) induced by interferon produced in response to infection of a small fraction of cells.

### ASTROCYTES ARE INNATE IMMUNE SENTINELS

To fully define ZIKV-regulated intracellular signaling pathways and their upstream regulators linked with DE gene expression, we used Ingenuity Pathway Analysis. This identified enrichment of interferon regulatory factor (IRF)3 and IRF7, occurring most strongly in astrocytes (Fig. 2A). IRF3 and IRF7 are critical transcription factors in signal transduction for innate immunity, linking the detection of viral RNA by pattern recognition receptors (PRRs) with innate immune activation and the downstream production of type I and III interferon (IFN) (Fig. 2B). We therefore compared baseline and induced expression of IRF3, IRF7, and additional genes marking innate immune activation and IFN response across cell types (Fig. 2C). Astrocytes demonstrated the highest expression levels of PRRs that are critical for ZIKV detection including retinoic acid inducible gene-I (RIG-I) and Toll like receptor (TLR)3^31,32^. These findings highlight a positive feedback loop engaged during viral infection, because both RIG-I and IRF7 are also canonical ISGs whose expression is induced by IFN following PRR signaling, and indicate that astrocytes are positioned to act as key innate immune sensors of ZIKV infection (Fig. S4D). Importantly, astrocytes expressed the highest levels of the type I interferon receptor (*IFNAR1* and *IFNAR2*, Fig. S4E).

**Figure 2.**
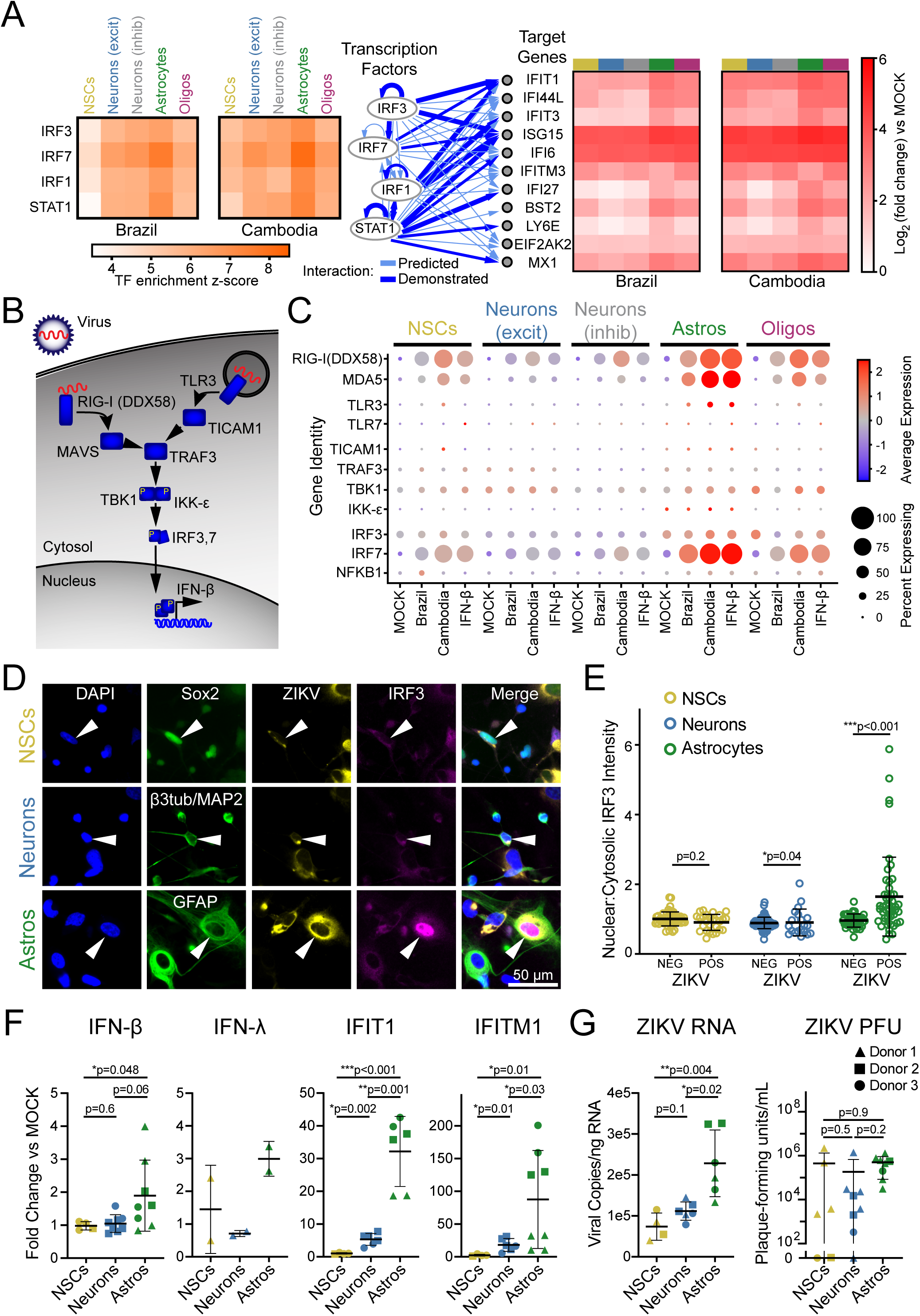
Astrocytes are innate immune sentinels in ZIKV infection. (A) Top four transcription factors (left) predicted by ingenuity pathway analysis to drive gene expression changes (right) observed in ZIKV infection. (B) Schematic indicating key steps in innate immune activation engaged by pattern recognition receptors hypothesized to detect ZIKV (RIG-I and TLR3). (C) Dot plot representing steady state, IFN-β-, and ZIKV-induced expression levels for innate immune signal transduction components. Color scale represents normalized expression (scaled median z-score across all genes and all conditions represented in the plot). (D) Immunocytochemistry (ICC) of innate immune activation, indicated by nuclear translocation of IRF3, in ZIKV-infected mixed human fetal primary brain cells (Donor 2). Cells were identified using the following ICC labeling: NSCs, Sox2 (top); neurons, combined β3-tubulin and MAP2 (middle); astrocytes, GFAP (bottom). (E) Quantification of IRF3 translocation, measured as the ratio of nuclear-to-cytosolic immunofluorescence, demonstrated innate immune activation only in ZIKV-positive astrocytes. N=1 experiment (3 coverslips quantified per cell type) from donor 2, representative of two similar experiments from two donors. Error bars reflect mean ± SD; p-value calculated by Kolmogorov-Smirnov test. (F) FACS-purified NSCs (gold), neurons (blue), and astrocytes (green) reflect cell type-specific response to ZIKV with astrocytes generating the strongest upregulation in IFN-β and the interferon-responsive genes IFIT1 and IFITM1. N=three donors, four independent experiments. (G) ZIKV RNA (left) and infectious titer (right) in the same conditions as F. Graphs in F-G demonstrate mean ± SEM. P-values represent unpaired t-test with Welch’s correction; one-way ANOVA p-value <0.05 for all graphs.

When activated by PRR stimulation, IRF3 and IRF7 form homo- or heterodimers and translocate to the nucleus to drive downstream gene expression, including IFNs. We therefore used immunofluorescence to analyze IRF3 subcellular localization during ZIKV infection in identified cell types (Fig. 2D). Astrocytes responded to ZIKV infection with nuclear translocation of IRF3, while NSCs and neurons did not (Fig. 2E). Astrocytes were also the only cell type that underwent ZIKV-driven nuclear translocation of NF-κB, which can also be induced by RIG-I and TLR3 signaling (Fig. S5A-B).

The relatively small proportion of astrocytes with nuclear IRF3 suggests that innate immune activation occurs sparsely even among infected cells. We surveyed the scRNAseq dataset to identify any cells containing reads for innate immune cytokines including type I, II, and III interferons as well as IL-6 and TNF-α (Fig. S5C). We found while very few cells expressed any of these cytokines, IFN-β was the sole type I interferon detected and was the only one expressed exclusively during ZIKV infection. Astrocytes were the predominant cell type expressing any cytokines in this analysis.

To confirm astrocytes are the source of ZIKV-induced IFN-β, we analyzed gene expression in specified cell populations purified from fetal brain primary explants. We adapted a fluorescence-activated cell sorting (FACS) methodology developed in iPSC-derived neural cells^33^, to obtain enriched populations of NSCs, neurons, and astrocytes from three donors (Fig. S2D-F). Upon infection with ZIKV, purified astrocytes induced expression of IFN-β, IRF3-target genes, and ISGs including IFIT1 and IFITM1, while purified NSCs and neurons showed little or no innate immune activation (Fig. 2F). Astrocytes also had higher copies of the ZIKV genome than NSCs and neurons (Fig. 2G).

### ZIKV INFECTION ENGAGES CELL-AUTONOMOUS APOPTOSIS PATHWAYS

Given the broad expression of ISGs across fetal brain primary cultures in response to ZIKV infection (Fig. 1E), we sought to distinguish ZIKV-specific cellular perturbations from the background ISG response. Using scPathoQuant, we compared gene expression in cells containing ZIKV reads^25^ to bystander cells from the same condition but lacking ZIKV reads (Fig. 3A, Fig. S6). This analysis removes the contribution of paracrine signaling including IFN-b from the transcriptional response, allowing the identification of virus-stimulated genes (VSGs) that are regulated independent of the direct actions of IFN. The strongest DE gene in this analysis was *CCL5* (chemokine C-C motif ligand 5) which was significantly upregulated only in astrocytes and was induced by both ZIKV strains. Because *CCL5* is a canonical innate immune chemokine induced downstream of RIG-I activation, this finding supports the conclusion that astrocytes act as innate immune sentinels in congenital ZIKV infection^34^.

**Figure 3.**
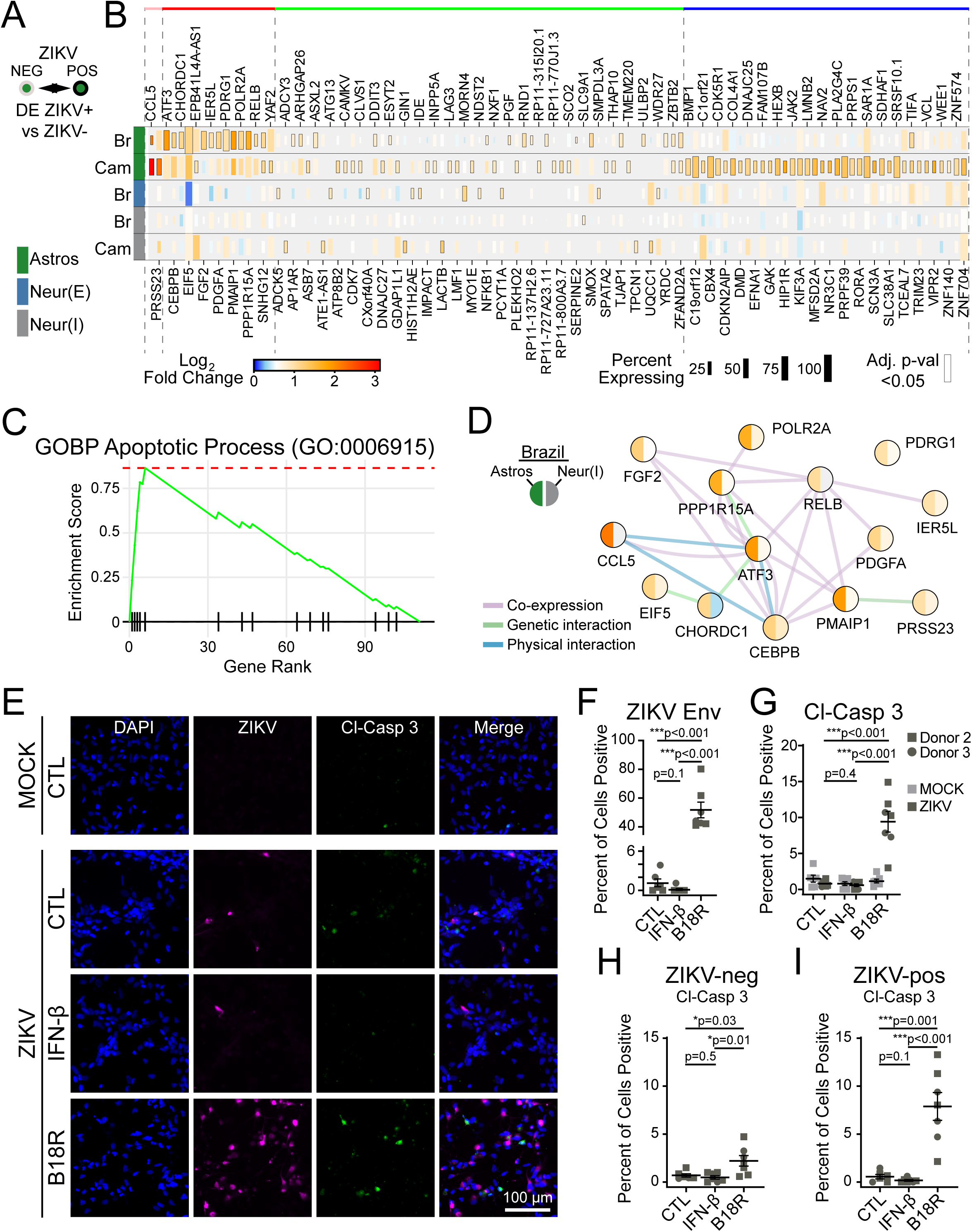
ZIKV infection drives cell-intrinsic apoptosis pathways limited by IFN-β. (A-B) Bioinformatic comparison (A), and heat map (B; rows=infection condition, grouped by cell type) identifying DE genes in cells containing ZIKV reads, compared to bystanders without ZIKV reads in the same treatment conditions. (C) GSEA identified apoptosis signaling as the top pathway induced in ZIKV-infected cells, LFCs were used to rank genes. (D) Network map of ZIKV-induced genes pertaining to intrinsic apoptosis (color=log-fold change by scale bar in B, according to ZIKV-Brazil condition). (E) Immunocytochemistry of cleaved caspase-3 (green) expression as a function of ZIKV-Brazil infection (magenta) in cells treated with IFN-β or the IFN-β inhibitor, B18R. (F-G) Blockade of IFN-β led to an increase in the percent of infected cells (F) and also the percent of cells staining positive for cleaved caspase-3 (G). (H-I) Percentage of cells with activated caspase-3 staining, when considering only ZIKV-negative (H) or ZIKV-positive cells (I). N=two independent experiments, two donors for panels F-I. All graphs indicate mean ±SEM.

We next used gene set enrichment analysis (GSEA) on the complete list of VSGs to identify key response pathways (Fig. 3B, Supplemental table 4). This analysis demonstrated strong enrichment for apoptosis signaling (GO:0006915) in ZIKV-infected cells compared to bystanders (Fig. 3C). This pathway includes the genes *PMAIP1, CEBPB*, and *ATF3* (Fig. 3C), which participate in caspase-mediated cell death linked to cellular stress^35–37^. Indeed, over-representation analysis (ORA) of ZIKV-induced genes identified additional signaling pathways linking intrinsic apoptosis to endoplasmic reticulum stress (GO: 0070059; GO: 0034976), particularly in astrocytes infected with ZIKV-Br (Fig. S6C). ZIKV, like related flaviviruses, extensively remodels the endoplasmic reticulum of host cells to form a replication compartment^38–40^, which can activate of cellular stress responses including the unfolded protein response (UPR)^41–43^. This observation may indicate a mechanistic link between ZIKV replication and apoptotic neural cell death.

To further explore apoptotic signaling in ZIKV-infected cells, we performed immunofluorescence analysis of cleaved caspase-3 in bulk fetal brain explant cultures (Fig. 3D). To achieve higher infection efficiency, we used vaccinia B18R protein, which selectively blocks type I interferon signaling^44,45^. Treatment with B18R led to an approximately 50-fold increase in the percent of cells infected with ZIKV in bulk cultures, highlighting the importance of IFN-β in restricting viral spread in fetal brain tissue (Fig. 3E-F). In addition, B18R increased the percent of cells staining for cleaved caspase-3 (Fig. 3E). Because B18R also increased percent of cells infected, we performed subgroup analysis of ZIKV-negative and ZIKV-positive cells (Fig. 3G-H). We found that ZIKV-infected cells treated with B18R were 13-times more likely to be positive for cleaved caspase-3 than ZIKV-infected cells treated with vehicle only.

Together, these findings indicate that ZIKV infection of human primary fetal brain tissue is detected primarily by astrocytes, leading to innate immune activation including production of IFN-β in a small fraction of cells that nonetheless causes widespread ISG induction in all cell types. Although ZIKV triggers apoptotic signaling pathways, IFN-β is protective against viral replication and apoptosis. Astrocytes therefore play a major role to impart innate immune protection against ZIKV infection and cell death in the fetal brain.

### HUMAN IPSC-DERIVED NSCS AND ASTROCYTES ARE HIGHLY SUSCEPTIBLE TO ZIKV INFECTION

Human primary fetal brain explants contained a small percentage of NSCs (2-5%), which limited our ability to explore ZIKV-related changes using scRNAseq. We therefore adapted a human cell model using induced pluripotent stem cells (IPSCs) to generate purified NSCs, applying a cell differentiation protocol to recapitulate the development of human frontal cortex^33^. This approach allowed us to produce NSCs and examine the consequences of ZIKV infection in undifferentiated NSCs (“STEM”) compared with mature neural cell types (“DIFF”, Fig. 4A, Fig. S7).

**Figure 4.**
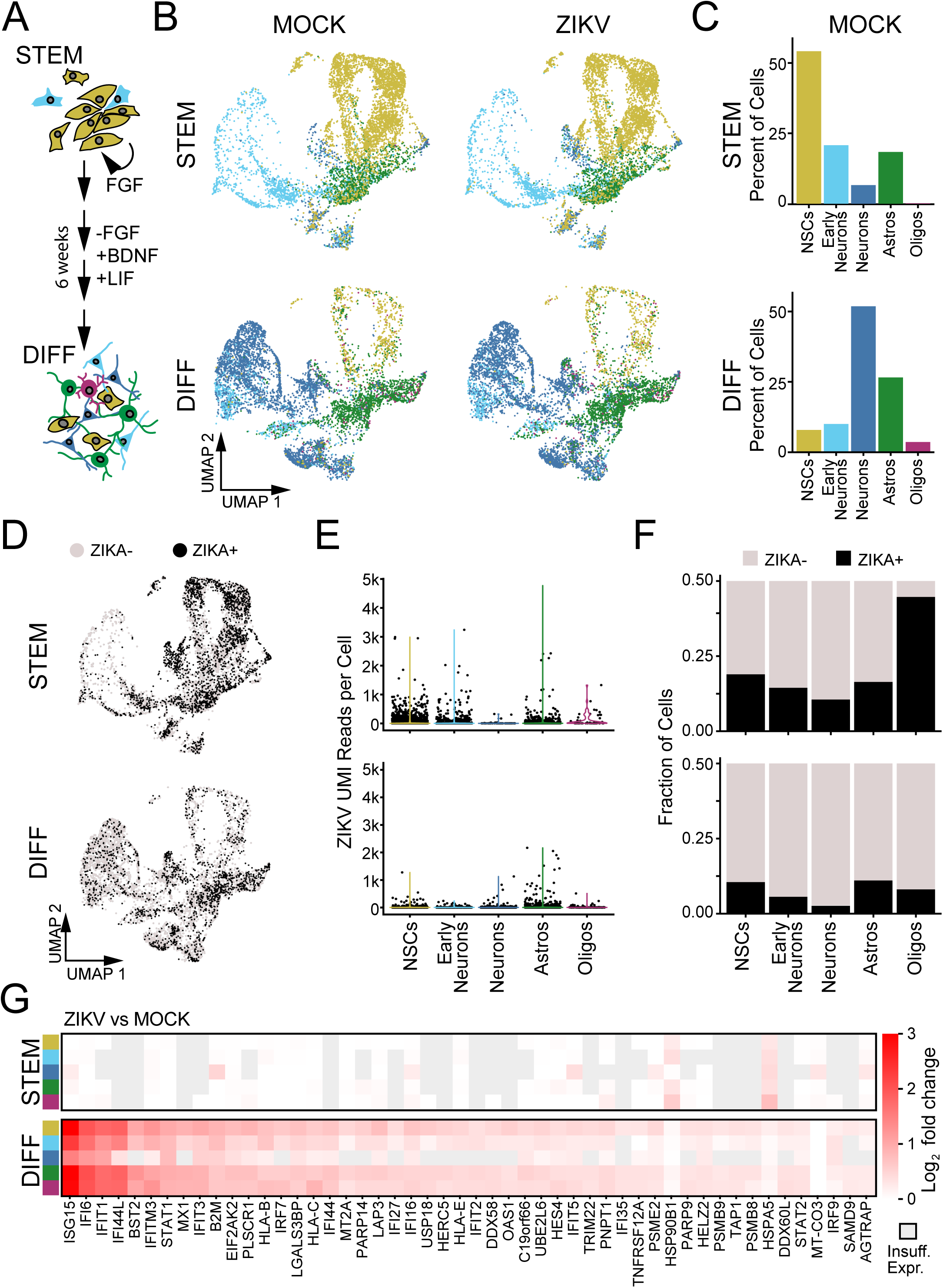
ZIKV replication and innate immune response in human IPSC-derived neural cells. (A) Schematic of differentiation protocol, which recapitulates development of human frontal cortex, beginning with purified neural stem cells (“STEM”) and resulting in mixed differentiated neural cell types (“DIFF”). (B) UMAP plots representing hIPSC-derived cells including neural progenitors (gold), early neurons (light blue), mature neurons (blue), astrocytes (green), and oligodendrocyte lineage cells (purple). (C) Quantification of the relative abundance (percentage) of each identified cell type in MOCK conditions. (D) UMAP plot of ZIKV-positive cells (black points, ≥9 ZIKV UMI reads/cell) and ZIKV-negative bystanders (see Fig. S9). (E) Violin plots of ZIKV UMI reads detected according to cell type (x-axis) in progenitors (top) and differentiated cells (bottom). (F) Percentage of ZIKV-positive cells by cell type and differentiation condition. (G) Heatmaps representing the top 50 DE genes (ZIKV vs MOCK “pseudobulk”) identified across cell types and differentiation states. The only significantly DE genes in progenitors (top) were HSP90B1 and HSPA5.

We performed scRNAseq after 48 hours of ZIKV infection in undifferentiated and differentiated cultures, using automated clustering to identify NSCs, immature (“early”) neurons, mature neurons (including inhibitory and excitatory), astrocytes, and oligodendrocytes (Fig. S8A). Undifferentiated cultures largely contained NSCs, with a small fraction of cells identified as early neurons and astrocytes. As the latter did not express protein markers of mature astrocytes (Fig. S7), they likely represent lineage-restricted progenitors. Differentiated cultures were predominantly composed of neurons. Astrocytes, oligodendrocytes, and NSCs were present in smaller proportions (Fig. 4B-C). This composition of cell types remained constant across ZIKV-infected and MOCK-infected cultures (Fig. S8B).

In IPSC-derived cells we detected an abundance of reads mapping to the ZIKV genome across all cell types (Fig. 4D-F), Fig. S9A). Based on sensitivity analysis in which we identified the ZIKV count threshold at which the maximal number of DE genes were identified across all cell types, we utilized a threshold of ≥9 ZIKV reads to define a cell as “virus-positive” (Fig. S9B-E). NSCs and astrocytes had the highest percentage of ZIKV-positive cells as well as the highest median counts of ZIKV reads (Fig. 4E-F). Similar to primary fetal brain explants, differentiated cultures demonstrated a robust transcriptional response to ZIKV infection featuring genes marking innate immune activation and the response to IFN-β (Fig. 4G, Fig. S10A, Supplemental Table 8). In stark contrast, undifferentiated NSCs did not exhibit induction of any ISGs and had almost no transcriptional response to ZIKV infection when examined at the population level (Fig. 4G). We again found that IFN-β was the predominant interferon subtype expressed during ZIKV infection, and ZIKV-positive astrocytes comprised the majority of cells expressing IFN-β transcripts (Fig. S10B). In contrast, undifferentiated cells did not contain any reads for IFN-β (Fig. S10B).

### ZIKV INDUCES PROGRAMS OF CELLULAR STRESS AND APOPTOSIS IN NSCS

To distinguish transcriptional changes induced by ZIKV from those driven by IFN-β in IPSC-derived cells, we performed separate DE analysis on ZIKV-positive and ZIKV-negative cells compared to MOCK infection for individual cell types (Fig. 5A-B). This analysis demonstrated ZIKV-driven gene changes according to cell type and differentiation state. For instance, one co-expressed gene module corresponding to the type I interferon response (top DE genes: *ISG15*, *IFI6*, *IFIT1*, *IFI44L*) was induced in all cell types regardless of ZIKV status but only in differentiated cultures (yellow module, Fig. 5B-C, Supplemental Table 9). This analysis also identified a “universal” ZIKV-related transcriptional response conserved across cell type and differentiation status but exclusively expressed in ZIKV-positive cells (red module, Fig. 5). Over-representation analysis (ORA) of this module demonstrated enrichment in pathways for cholesterol synthesis, cellular stress, and the unfolded protein response (Supplemental Table 10).

**Figure 5.**
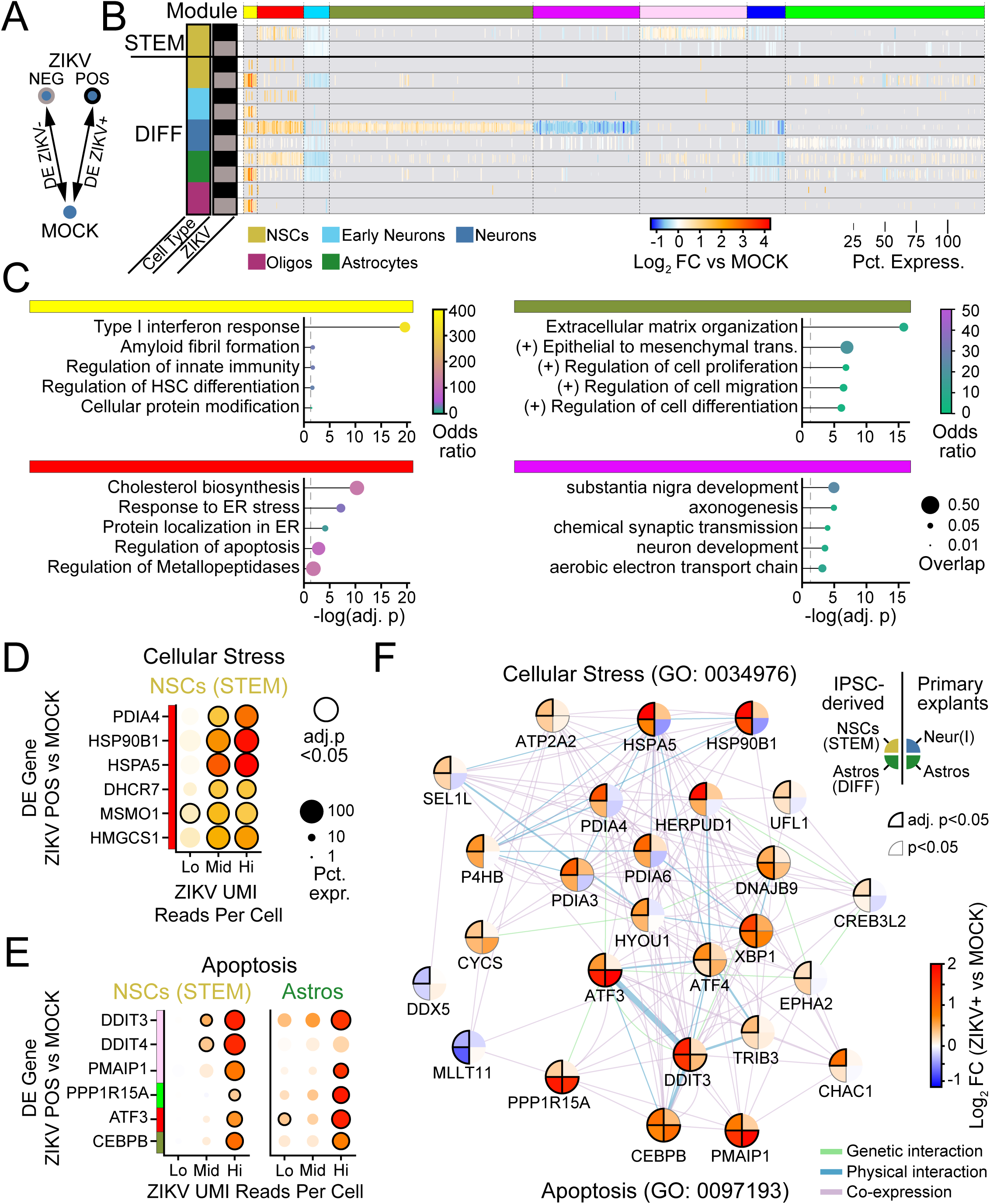
Universal ZIKV-driven transcriptional responses reflect cellular stress linked to apoptosis. (A) Schematic of DE comparison. (B) Heatmap of DE genes according to cell type (row color) and ZIKV-positivity (grey and black row markers). K-means clustering of genes identified co-expression modules (top color bar) that are universal across cell types (e.g., red), modules that are specific to differentiated cells (e.g. yellow), and genes that are specific to neurons (magenta and forest green modules). (C) GSEA of four modules from B indicates that universal gene modules include IFN-β signaling (yellow) and cellular stress (red), while neuron-specific gene modules include extracellular matrix remodeling (forest green) and synaptic function (magenta). (D-E) Dot plots representing gene expression changes in ZIKV-positive cells (vs MOCK) as a function of the number of ZIKV UMI reads (x-axis), for genes pertaining to cellular stress (D) or apoptosis (E). Lo: 1-9 reads; Mid: 10-99 reads; Hi:>99 reads. Color: log-fold change indicated by color bar in F. (F) Network map linking cellular stress and apoptotic signaling in ZIKV-infected cells for iPSC-derived cells (left half) and primary fetal explants (right half). Color: log-fold change versus MOCK (for iPSC-derived cells, from cells with >99 ZIKV UMI reads).

The abundance of viral reads in IPSC-derived cells allowed us to perform transcriptional analysis as a function of viral load. To do so, we subdivided ZIKV-positive cells into those with low (<10), middle (10-99), and high (≥100) read counts. We used this approach to test whether ER cellular stress pathways might provide a link between viral replication and cell death via apoptosis that could be sensitive to viral load. Indeed, this analysis approach demonstrated that key genes representing cellular stress (*PDIA4, HSP90B1*, *HSPA5*) and contributing to apoptosis (*DDIT3, PMAIP, ATF3*) were induced to higher levels with increasing ZIKV copy number (Fig. 5D-E).

To interrogate mechanistic links between these pathways, we defined genes comprising endoplasmic reticulum stress (GO: 0034976) and intrinsic apoptosis gene networks (GO:0097193; Fig. 5F) that were significantly perturbed in ZIKV-infected cells. Although genes in these pathways were identified in IPSC-derived cells, many genes overlapped with the apoptosis pathways identified in primary human fetal explants (for example see Fig. 3D), allowing us to extend and verify our analysis of ZIKV-induced cell death signaling pathways. We found widespread correlation in ZIKV-induced gene expression in cells across the tissue types, particularly in genes associated with intrinsic apoptosis (*ATF3*, *DDIT3*, *PPP1R15A*, *CEBPB*, *PMAIP1*). Together, these findings identify a universal gene signature of ZIKV infection in neural cells, in which viral load-dependent ER stress gene induction links with expression of apoptotic cell death genes regardless of cell type.

### ZIKV PERTURBS DEVELOPMENTAL PROGRAMS IN NEURONS

Analysis of ZIKV-stratified DE genes also identified co-expression modules that were unique to specific neural cell types (Fig. 5B). We found that neurons had the highest number of ZIKV-regulated genes in that 483/1345 (35%) of all DE genes were specific to neurons, and that neurons had the largest magnitude ZIKV-related changes in gene expression among cell types (Fig. 6A-B).

**Figure 6.**
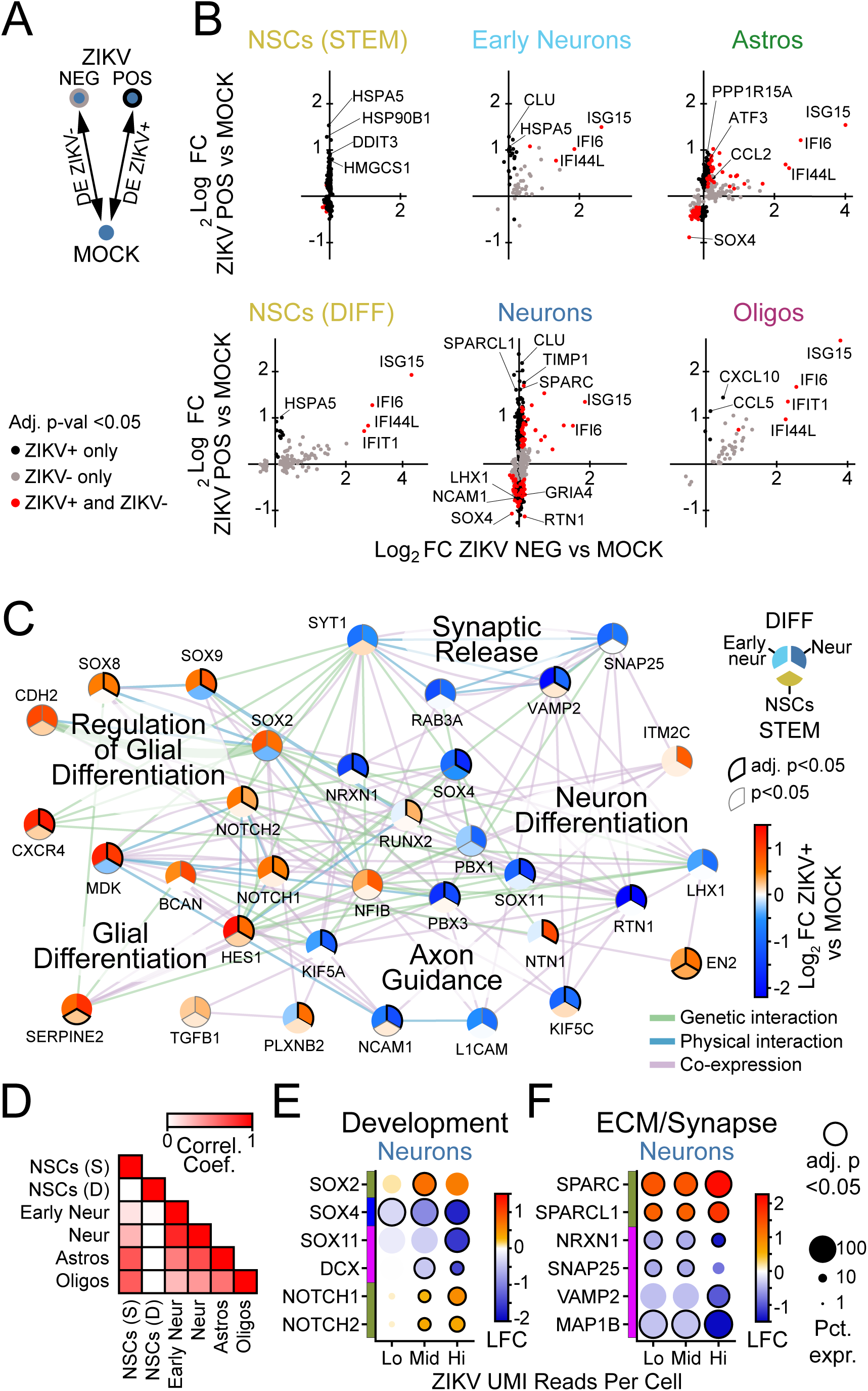
Pathways of neurodevelopment and synaptic function are perturbed in ZIKV-infected neurons. (A) Schematic of DE comparison. (B) Scatterplots (points=genes) representing log-fold change in ZIKV-positive cells (x-axis) versus ZIKV-negative cells (y-axis). Color indicates conditions in which significance (adj. p-val. <0.05) was reached. (C) Network of gene expression programs perturbed in ZIKV-infected neurons (upper right third), early neurons (upper left third), and NSCs (lower third). Color=log-fold change in cells with >99 ZIKV UMI reads versus MOCK, per color bar. (D) Correlation matrix indicating correlation coefficient calculated using log-fold change (ZIKV-positive vs MOCK) of genes shown in (C), in pairwise comparison between cell types as indicated on x and y axis. (E-F) Dot plots representing gene expression changes in ZIKV-positive cells (vs MOCK) as a function of the number of ZIKV UMI reads (x-axis), for genes pertaining to synapses (D) or neuron development (E) in neurons, or innate immunity (F) in astrocytes. Lo: 1-9 reads; Mid: 10-99 reads; Hi:>99 reads. Color: log-fold change indicated by color bar in F.

ZIKV-infected neurons demonstrated virus-induced gene modules related to extracellular matrix remodeling and virus-suppressed gene modules related to neural development (Fig. 5C, forest green and magenta modules, respectively). To explore how ZIKV infection may affect neural development, we performed ORA of all DE genes in ZIKV-positive neurons (Supplemental table 10) from which we selected genes from GO terms relating to neuronal differentiation (GO:0030182) and function (GO:0099643 and 0007411), and glial differentiation (GO:0045687 and 0010001) for further interrogation. A network map of these related genes demonstrated a global shift or “switch” in ZIKV-infected neurons away from transcriptional patterns associated with neuron-specialized functionality and toward transcriptional patterns associated with glial cell development (Fig. 6C).

The direction and magnitude of ZIKV-related gene expression changes in this network analysis were strongly correlated between neurons and early neurons, while less so with other cell types (Fig. 6D). In particular, NSCs did not demonstrate perturbations in pathways for neuronal function to favor pathways of glial differentiation, highlighting the specificity of ZIKV-dependent perturbations of differentiation for cells in the neuronal lineage. At increasing levels of ZIKV viral RNA, expression levels of genes related to glial development and ECM remodeling increased, while expression of genes for synaptic function decreased (Fig. 6E-F), indicating that the observed perturbations in neuron signaling pathways were closely linked to ZIKV replication. We did not observe a spatial concentration of ZIKV-positive cells within any cluster of mature neurons in UMAP space, arguing against a particular subtype of neurons having increased susceptibility to ZIKV infection (Fig. S8C).

### NSCS DO NOT MOUNT AN INNATE IMMUNE RESPONSE TO ZIKV INFECTION

Finally, we further examined NSC gene expression by comparing bulk cultures of IPSC-derived neural cells before and after differentiation. In contrast to differentiated neural cells, NSCs did not express genes marking innate immune activation (see Fig. 6B). We found that, in response to ZIKV infection, cultures of differentiated neural cells undergo phosphorylation of TBK1, IRF3, and STAT1 but undifferentiated NSCs do not, suggesting that they do not sense virus infection (Fig. 7A). Single cell analysis of specific PRR expression revealed low basal RIG-I mRNA expression in undifferentiated NSCs but increased basal RIG-I mRNA expression in differentiated NSCs similar to levels in glial cells (Fig 7B). RIG-I levels were further induced following IFN-β treatment, similar to that observed in primary fetal explants (see Fig. 2C, Fig. S10C). TLR3 expression was low in all cells and was only minimally increased in differentiated astrocytes after IFN-β treatment. Moreover, immunoblot analysis showed that undifferentiated NSCs had remarkably low levels of RIG-I protein compared to abundant RIG-I protein in differentiated neural cell cultures (Fig.S7C). By comparison, TLR3 protein expression was highest in undifferentiated NSCs. To clarify innate immune sensing capacity, we assessed the response to treatment of bulk cultures with RIG-I agonist (poly-U/UC 5’ppp RNA)^46^ or TLR3 agonist poly-I:C RNA. We observed dose-dependent IFN-β production in response to either agonist in differentiated neural cultures but not in undifferentiated NSC cultures (Fig. 7C-D).

**Figure 7.**
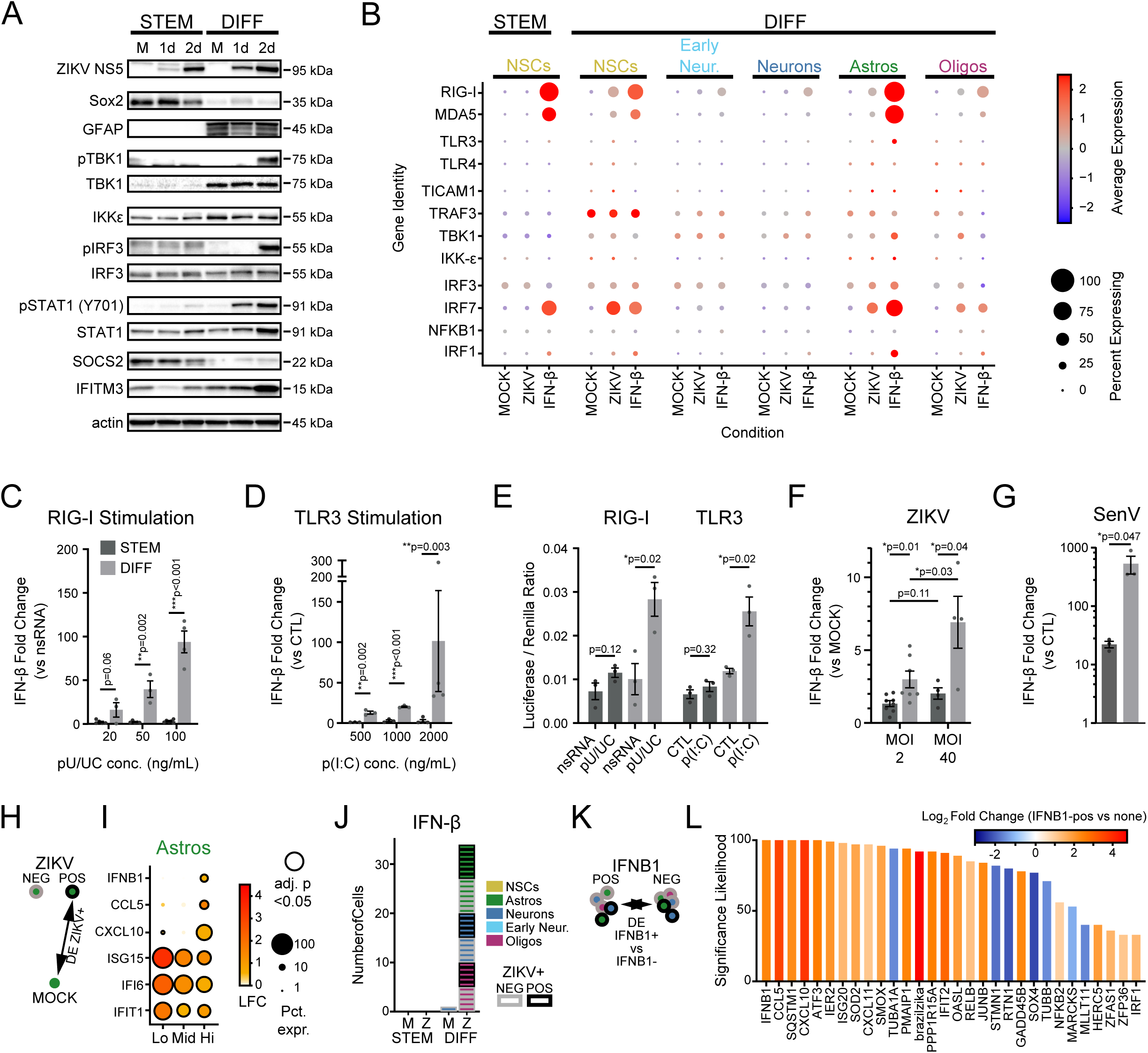
Failure of innate immune signaling in neural stem cells. (A) Innate immune activity in neural stem cell (left) and differentiated cultures (right), as measured by phosphorylation of key signaling proteins (TBK1, IRF3, STAT1) after 1 and 2 days of ZIKV infection in bulk cultures. (B) Dot plot representing steady state, IFN-β-, and ZIKV-induced expression levels for innate immune signal transduction components in scRNAseq data. Color scale represents normalized expression (scaled median z-score across all genes and conditions in the plot). (C-D) IFN-β gene induction in bulk cultures by qPCR in neural stem cell (dark grey) and differentiated cultures (light grey), induced by treatment with a RIG-I agonist (C, poly(U/UC) transfection) or TLR3 agonist (D, poly(I:C) extracellular application). (E) Firefly luciferase activity, induced under control of an exogenous IFN-β promoter in response to poly(U/UC) or poly(I:C). Firefly luciferase was normalized to Renilla luciferase driven by a CMV promoter to control for transfection efficiency. (F-G) IFN-β induction in response to high ZIKV inoculum or the strong RIG-I agonist, Sendai virus. P-values on graphs in C-D represent unpaired t-test. (H-I) Differential gene expression comparing ZIKV+ cells to MOCK according to level of ZIKV RNA. Astrocytes significantly upregulated innate immune signaling genes (CXCL10, CCL5, IFNB1) at the highest level of ZIKV reads. (J) IFNB1-expressing cells according to cell type (color) and ZIKV infection status (border) across treatment conditions. (K-L) Differentially expressed genes (x-axis) in IFNB1-expressing cells as identified in (J), compared to IFNB-negative cells in the same treatment condition. Significance likelihood reflects bootstrap analysis of 100 DE comparisons and is a measurement of the percent of comparisons in which the gene fold change was identified as significant with adjusted p-value <0.05. Color reflects average log_2_-fold change across all comparisons.

In some tissues, pluripotent stem cells are incapable of IRF3/7-driven IFN-β production, due to chromatin inaccessibility of the promoter region^10,47^. To evaluate whether this mechanism persists in neurally-restricted progenitors, we transfected cells with a plasmid containing a luciferase gene under control of an exogenous IFN-β promoter that would not be subject to expression regulation by chromatin modification. Differentiated cells responded to RIG-I or TLR3 agonists with robust luciferase production, while undifferentiated NSCs did not (Fig. 7E). While infecting NSCs with ZIKV at high multiplicity of infection did not engage innate immune signaling, infection with Sendai virus (which strongly activates RIG-I) did lead to a weak induction of IFN-β, albeit far smaller than that observed in differentiated cells (Fig. 7F-G).

We next sought to identify transcriptional signatures corresponding to innate immune activation in differentiated cells using the total scRNAseq dataset. In astrocytes with the highest ZIKV copy number, this analysis identified induction of innate immune genes including IRF3-target genes and ISGs marked by *IFIT1, ISG15, IFNB1*, *CXCL10*, and *CCL5* (Fig. 7H-I). This gene expression signature indicates that, as in human primary fetal explants, iPSC-derived astrocytes act as innate immune sentinels, and they support high levels of ZIKV replication while producing IFN-β. We also noted a trend in the expression of ISGs in which high viral load linked with a reduction of ISG expression, consistent with ZIKV-mediated suppression of interferon signaling^48^ (Fig. 7I)

IPSC-derived neural cells demonstrated ZIKV-related IFN-β production in neurons and oligodendrocytes, although not in NSCs (Fig 7J). To define gene expression profiles that might underlie a cell’s capacity to engage innate immune signaling, we performed DE analysis comparing cells containing *IFNB1* reads to cells from the same culture condition that lacked *IFNB1* expression (Fig. 7K). By this analysis, *IFNB1*-positive cells were marked by higher levels of *CCL5, IFIT2,* and *CXCL10*, indicating these cells are undergoing innate immune activation. In contrast, we found decreased levels of *SOX4* in *IFNB1*-positive cells (Fig. 7L). Because Sox4 inhibits innate immune signal transduction at multiple levels^49,50^, this may be a mechanism contributing to the failure of NSCs to induce IFN-β in response to ZIKV infection. Together, these data indicate that impaired innate immunity in NSCs likely occurs at multiple levels, including decreased expression levels of PRRs, failure to engage phosphorylation of downstream signaling partners, and possible inhibition by Sox4.

## Discussion

The 2016 outbreak of ZIKV in Brazil alerted the world to a new etiology of congenital infection with devastating neurologic consequences. Although CZS outcomes are now understood to range from subtle developmental delay to severe microcephaly, the mechanisms of neurologic injury in CZS remain poorly understood^51–53^. We used scRNAseq in human primary brain explants and IPSC-derived neural cells to achieve holistic insight into the consequences of ZIKV infection across diverse fetal brain cell types for the first time, developing the Human Neural Cell Atlas of ZIKV Infection. Importantly, the Atlas is inclusive of ZIKV RNA reads, allowing us to distinguish virus-induced (cell-intrinsic) from virus-independent (paracrine) transcriptional responses. This approach provides mechanistic insight into human fetal brain neural cell gene expression and regulation by ZIKV in three areas: innate immune function, apoptosis, and neurodevelopment, that should serve to inform our understanding of congenital ZIKV infection and CZS.

ZIKV had broad tropism across cell types spanning both primary and IPSC-derived tissue, in agreement with studies that have described ZIKV replication in neural cells other than NSCs^17,19,20,54^. We found that innate immune signaling was engaged principally in astrocytes and propagated by IFN-β, which dominated the transcriptional response to ZIKV in all neural cells examined. Strikingly, we found that NSCs failed to induce IFN-β in response to ZIKV infection, although they upregulated ISGs in response to IFN-β. While some studies of ZIKV infection have identified functional innate immunity in NSCs^12–14^, others have not^7,15^. We propose these disparate results fundamentally reflect the differentiation state of the progenitor populations in the culture conditions being studied.

Both primary and embryonic stem (ES) cell-derived NSCs undergo rapid initial differentiation in 2D culture and organoids, and it is difficult to retain purity of progenitor populations^55,56^. In contrast, IPSC-derived cells are thought to have a slower time course for differentiation^57^. Notably, one study examining ZIKV infection of ES cell-derived neural progenitors reported the magnitude of the innate immune response increased as cells differentiated^13^. To identify NSCs using immunofluorescence, we used Sox2 as a marker, which has been linked to ZIKV susceptibility^58^. We also identified *SOX4*—a closely related transcription factor that is expressed in immature neurons—as inversely correlated with IFN-β production in ZIKV-infected cells. By comparing multiple cell types and stages of differentiation simultaneously, our approach situates innate immunity of primary and IPSC-derived NSCs in a developmental context, arguing they have lost intrinsic immunity of pluripotent stem cells^9^ but have not yet acquired innate immune sensing capacity. Indeed, our findings in IPSC-derived NSCs suggest a multi-level impairment in innate immune signaling, including lower expression of RIG-I and failure to activate IRF3 (Fig. 5). Viral strain also influences innate immune responses, Asian-lineage ZIKV isolates induce less IFN-β than ancestral African-lineage strains^59^. We found that the epidemic Brazilian strain produced a weaker host transcriptional response than the older Asian-lineage strain, as we have reported previously^30^. Therefore, viral and host properties may interact to blunt the innate immune response in NSCs.

One notable limitation of our study was the availability of gestational ages for primary tissue (12-17 gestational weeks [gw]), which allowed us to capture key neural cell types but precluded analysis of events earlier or later in neurodevelopment. To identify NSCs within our scRNAseq data sets, we used *PAX6* and *PCNA* as NSC markers, which are highly expressed in ventricular zone progenitors of first trimester fetal brain^60^. Fetal brain development proceeds through an initial phase of stem cell expansion during the first trimester, followed by phases of predominant neurogenesis and then astrogliogenesis. Notably, the incidence of microcephaly is highest when initial ZIKV infection occurs during the late first trimester (10-14gw)^61^, corresponding to a developmental time point at which post-mitotic neurons are just beginning to populate the brain^62^, while mature astrocytes appear after 14gw^63^. As our observations show that astrocytes are a cell type sensing and responding to ZIKV infection to produce IFN-□, ZIKV infection prior to astrocyte differentiation of NSCs would confer high susceptibility of NSCs to ZIKV infection. Indeed, we observed dramatic protection conferred by IFN-β in fetal brain primary cells against ZIKV infection and apoptotic signaling of cell death. The high susceptibility of NSCs links with their low level of RIG-I expression, the key PRR responsible for sensing ZIKV and triggering the innate immune response to infection^30^, and reflects their lack of response to ZIKV and RIG-I agonist RNA treatment. Thus, impaired innate immunity in NSCs due to low RIG-I expression contributes to their increased susceptibility to ZIKV. ZIKV infection of NSCs and other neural cells can be suppressed by the actions of IFN produced by astrocytes but this protection is likely to be temporally and spatially restricted.

We have recently described methods for the simultaneous identification and alignment of virus and host reads in single cell RNA transcription datasets^25^. Here, we integrate that methodology to correlate levels of viral replication with host gene expression. Output from this approach revealed gene expression modules constituting the cellular response to endoplasmic reticulum stress induced across neural cell types as a function of ZIKV RNA levels (see Figs. 3,5). Flaviviruses including ZIKV replicate in the endoplasmic reticulum compartment and trigger the unfolded protein response^42,43^, an outcome that supports the validity of our virus-inclusive scRNAseq approach. Our bioinformatics analysis extended this module to link ER stress to cell death via pathways underlying intrinsic apoptosis (Figs. 3,6), and our findings are corroborated by several prior studies to the level of specific genes including *PMAIP* and *CEBPB*^64,65^. Despite activation of intrinsic apoptosis pathways, we did not observe extensive cell death in ZIKV-infected neural cells when innate immune signaling was intact. However, IFN-β blockade led to a significant increase in activated caspase-3 (Fig. 3), indicating innate immune signaling limits cell death in ZIKV infection of neural cells. Our findings suggest ISGs exert anti-apoptotic effects by suppressing ZIKV infection and replication to limit ER stress and engagement of the unfolded protein response. However, we cannot formally exclude the possibility that one or more ISGs exert a direct anti-apoptotic effect, as has been described for IFI6^66^.

Several studies investigating ZIKV-related perturbations in neurodevelopment have proposed that ZIKV limits NSC proliferative capacity^7,8^ or decreases generation of neurons^5,14,15^. Importantly, our data suggest ZIKV-linked perturbation in neurodevelopmental pathways occurs not in NSCs but rather in cells restricted to the neuronal lineage (see Fig. 7). Across all cell types, IPSC-derived neurons had the largest number of ZIKV-induced transcriptional changes. ZIKV infected neurons had increased expression of several genes that contribute to extracellular matrix (ECM) modification, including *TIMP1, SPARC,* and *SPARCL1*. During neurodevelopment, ECM remodeling is a key mechanism for controlling synapse number^67^, and the secreted protein encoded by the *SPARC* gene acts *in vitro* in a dose-dependent fashion to cause synaptic disconnection, axon retraction, and decreased expression of neurotransmitter receptors^68–70^. We also found that genes responsible for neurite outgrowth (*TUBA1A*, *NCAM1*, *MAP2*, *STMN1/2*) and synaptic function (*NSG1*, *NRXN1*, *NRXN2*) were strongly suppressed in ZIKV-infected neurons, while genes directing glial cell lineage differentiation (*SOX8*, *SOX9*, *HES1*) were induced by ZIKV in neurons. We propose this functional dichotomy in gene neuronal gene expression represents a “switch” from neurogenesis to gliogenesis induced by ZIKV infection. Gliosis is a hallmark pathologic feature of the ZIKV-infected fetal brain^27,28^ and is demonstrated in nonhuman primate models of maternal-fetal ZIKV infection^29,71^. The Human Neural Cell Atlas of ZIKV Infection now reveals the gene signature of this neuronal to gliosis switch, allowing full interrogation of involved gene expression to enable a molecular definition of ZIKV pathogenesis.

Together, these findings suggest that ZIKV infection in neurons may engage developmental programs that lead to de-differentiation and even to synapse elimination. Other data indicate ZIKV infection leads to a decrease in synaptic gene expression in a non-human primate model of congenital ZIKV infection^72^ as well as mouse^73^ and tissue culture models^54^ of ZIKV infection. One aspect of CZS that is poorly understood is the progression of microcephaly late in gestation or even after birth. This process has been noted to resemble Fetal Brain Disruption Sequence, in which death of post mitotic neurons is thought to the central mechanism, as opposed to loss of progenitors or decreased neurogenic output^1^. Newborn cortical neurons are dependent on the formation of functional synapses for survival and an estimated 50% of post mitotic neurons undergo programmed cell death during development as neural circuits are refined^74^. Our data suggest that late development of microcephaly may reflect neuronal apoptosis resulting from insufficient synapse formation and trophic support, in addition to direct virus-driven expression of apoptotic gene networks identified in this study.

The Human Neural Cell Atlas of ZIKV Infection provides significant insight and a new tool for interrogating the mechanisms of neuropathology in congenital ZIKV infection of fetal neural cells at the single cell level. The Atlas reveals diverse cellular responses reflecting distinct innate immune- and virus-driven pathways and their dynamic interplay. We show that type I interferon signaling, driven by astrocyte-derived IFN-β, is a key determinant of the cellular response to ZIKV infection. We propose that impaired innate immunity of NSCs is due to their expression of only low levels of RIG-I to underlie their unique high susceptibility to ZIKV infection, allowing for high levels of viral replication, progression of disease, and CZS. We also identify cell-intrinsic programs of virus-driven gene expression, including neuron-specific responses that may lead to loss of synapses and de-differentiation of neurons, with widespread ramifications for neurodevelopment. Finally, we identify innate immune, apoptotic, and differentiation pathways that may represent therapeutic targets for preventing the most severe consequences of CZS.

## Methods

### RESOURCE AVAILABILITY

#### Lead contact

Further information and requests for resources and reagents should be directed to and will be fulfilled by the lead contact, Michael Gale, Jr.

#### Materials availability

This study did not generate new unique reagents.

#### Data and code availability

Single-cell RNA-seq data have been deposited in the Gene Expression Omnibus (GEO) and are publicly available as of the date of publication. Accession numbers are listed in the key resources table. Original microscopy and western blot images reported in this paper will be shared by the lead contact upon request.

All original code will be deposited at the Gale lab site on github, https://github.com/galelab/Stokes_SC_Atlas_ZIKV_brain_infection.git and is publicly available as of the date of publication. DOIs are listed in the key resources table. Experimental treatment conditions are included in the GEO repository and described in detail below; any additional information required to reanalyze the data reported in this paper is available from the lead contact upon request.

### EXPERIMENTAL MODEL AND STUDY PARTICIPANT DETAILS

#### Human primary fetal brain tissue

De-identified human primary fetal (12-17 weeks post-conception) brain tissue was obtained from the Birth Defects Research Laboratory (BDRL) at the University of Washington with ethics board approval and maternal written consent. This study was performed in accordance with ethical and legal guidelines of University of Washington Institutional Review Board.

Gross dissection was performed by BDRL staff to isolate developing forebrain, tissue was stored in Hibernate E medium at 4°C (Gibco) for transport, then processed within 24 hours. For immunohistochemistry, tissue was immediately fixed in 4% paraformaldehyde at 37°C. For cell culture, tissue was dissected into 1mm^3^ blocks using a scalpel, then transferred into 4°C Accutase digestion medium (Invitrogen) for one hour, with gentle trituration every 10-15 minutes through progressively smaller bore pipettes. Once a single cell suspension was obtained, this was plated as a monolayer.

Human primary fetal cells were maintained at 37°C in a 5% CO2 incubator on tissue-culture grade plastic dishes or glass coverslips (Carolina Glass, #633029) coated with 20 μg/mL poly-L-ornithine (Sigma, P-3655) and 10 μg/mL laminin (Gibco) for four hours at 37°C. Human primary fetal tissue was maintained in differentiation medium (DD) consisting of DMEM/F12 (Gibco #11320033) base, containing: 1x B27 without Vitamin A (Gibco #12587010), 1x N-2 (Gibco #17502048), 1x antibiotic/antimycotic (Gibco #15240062), and 1mM L-glutamine (Gibco #25030081), supplemented with 20 ng/mL brain-derived neurotrophic factor (BNDF; Peprotech #450-02) and 20 ng/mL leukemia inhibitory factory (LIF; Peprotech #300-05). Cells were fed every 2-3 days with fresh medium.

Freshly obtained primary cells were recovered and expanded for 7-14 days on 15cm dishes, then passaged using Accutase digestion and plated onto 6-well plates (5e5 cells per well) for scRNAseq and Western Blot experiments or 12-well plates (2.5e5 cells per well) for qPCR experiments. Human primary cell cultures were treated empirically for Mycoplasma with 25 mg/mL plasmocin (InvivoGen #ant-mpt-1) after initial isolation for a minimum of two weeks unless used before this for experiments. Human primary fetal tissue was only used after it was confirmed negative for mycoplasma contamination using the MycoAlert Mycoplasma test kit from Lonza (#LT07-318).

#### Human induced pluripotent stem cells

Human induced pluripotent stem cells (hIPSCs, cell line CVI-A2; male) were differentiated into neural progenitors using dual-SMAD inhibition techniques as previously described^33,75^. NSCs were maintained on 10cm dishes in growth medium (GM) consisting of DMEM/F12 (Gibco #11320033) base, containing: 1x B27 (Gibco #17504044), 1x N-2 (Gibco #17502048), 1x antibiotic/antimycotic (Gibco #15240062), and 1mM L-glutamine (Gibco #25030081), supplemented with 20 ng/mL basic fibroblast growth factor (bFGF; Sigma-Aldrich #GF003). NSCs were passaged using Accutase and plated onto 6-well plates (3e5 cells per well) for scRNAseq and Western Blot experiments, 12-well plates (2e5 cells per well) for qPCR experiments, or 24-well plates (5e4 cells per well) for immunofluorescence experiments. Differentiation into mixed neural cells was performed by switching NSCs from GM medium to DD medium for a period of 6 weeks prior to experiments. To control for alterations in signaling due to treatment medium, all iPSC-derived cell types (both NSCs and differentiated cells) were switched to GM medium, supplemented with 20 ng/mL BDNF and 20 ng/mL LIF for 48 hours prior to infection or other treatment. All iPSC-derived cell lines were tested and negative for mycoplasma contamination using the MycoAlert Mycoplasma test kit from Lonza (#LT07-318).

### VIRUS STOCKS AND INFECTION

The following viruses were used in this study: Zika virus epidemic strain isolated in the Brazilian state of Paraiba in 2015 (Genbank KX811222; termed ZIKV Brazil); Zika virus strain FSS13025 isolated in Cambodia in 2010 (Genbank KU955593; termed ZIKV Cambodia)^30^. Working stocks of virus were generated from plaque-purified isolates, and amplified in Vero E6 cells, which were maintained in Dulbecco’s modified Eagle medium (DMEM) supplemented with 10% fetal calf serum (Cytiva #SH3007103), 10 mM L-glutamine, 1 mM sodium pyruvate (Corning #MT25000CI), 1x antibiotic-antimycotic solution, and 10 mM HEPES (Corning #MT25060CI). Virus stock titers were obtained using plaque assay on Vero cells and also by focus-forming assay on iPSC-derived NSCs, as previously described^30^. Multiplicity of infection (MOI) is reported based on Vero-derived titers. All virus stocks were sequenced to confirm identity, using QIAmp viral RNA mini kit (Qiagen #52906) for RNA extraction followed by library preparation using the KAPA RNA HyperPrep Kit with RiboErase (HMR) (Roche Diagnostics) and next-generation sequencing on the Illumina NextSeq 500 Instrument using a High Output 150 cycle kit. For single cell RNA-sequencing experiments in iPSC-derived tissues, to achieve higher multiplicity of infection, the ZIKV Brazil isolate was amplified in NSCs maintained in GM medium which does not contain serum. The resulting viral stocks had 99.9% sequence identity by nucleotide with the parental Brazil strain. Phenotypic consequences including kinetics of replication and transcriptional response of NSC-grown viral stocks were indistinguishable from Vero-grown viral stocks. All viral stocks were regularly tested and negative for mycoplasma.

Sendai Virus (Cantell strain, ATCC VR-907) infection was performed as previously described^30^, with minor modifications as follows. Adsorption of 100 HAU/mL was performed in low volume on a rocking platform at 37°C for 6 hours, after which a full exchange of medium was performed.

### SINGLE CELL RNA SEQUENCING

Single-cell 3’ RNA sequencing was performed according to the specifications of the manufacturer (10x Genomics). Briefly, cells were grown as above at a density of approximately 7.5e5 cells per well in a 6-well plate, then released from the dish using Accutase (Invitrogen, #00-4555-56), and triturated to generate a single cell suspension. Cells were loaded with viability dye (Invitrogen #65-0865-14) and sorted on a FACSAria II cell sorter (BD Biosciences) to isolate live cells. This single-cell suspension was then loaded onto a Chromium Single Cell Chip G to produce gel beads-in emulsion (GEMs) at a target capture rate of 5,000 cells per sample (MOCK and IFN-β treated) or 8,000 cells per sample (ZIKV-infected). All samples from a single experiment were loaded simultaneously to reduce variability. Barcoded, full-length cDNAs were produced by reverse transcription, and indexed libraries were prepared from 10 µl of cDNA using Chromium Next GEM Single Cell 3’ Reagent Kits v3 (10x Genomics). Library and cDNA quality were evaluated using a Bioanalyzer 2100 Instrument (Agilent) and quantifed using a ViiA 7 Real-Time PCR system (TermoFisher). Constructed libraries were sequenced on a NovaSeq 6000 (Illumina). Each sample was sequenced at a targeted read depth of 80,000 reads per cell. This required a second sequencing run for the human primary fetal tissue, achieving a final read of 120,000 reads per cell.

### BIOINFORMATICS ANALYSIS OF SINGLE CELL TRANSCRIPTIONAL DATA

Single cell reads were aligned to the hg19 genome with CellRanger software (10x Genomics; v5.0.1). Remaining reads were aligned to the respective ZIKV genomes for Brazilian (<u>KX811222</u>.1) and Cambodian (MH158236.1) isolates using scPathoQuant^25^. Seurat software (www.satijalab.org/seurat/; v4) was used to normalize gene expression, perform dimensionality reduction for principle component analysis (PCA) followed by uniform manifold projection (UMAP), cell clustering, and identify differentially expressed genes. Initial quality control included filtering out cells that had less than 200, a mitochondrial DNA percentage greater than 10% and those that were predicted to be doublets using DoubletFinder (https://www.sciencedirect.com/science/article/pii/S2405471219300730). For cell type identification we used Garnett software^76^ (cole-trapnell-lab.github.io/garnett/; v0.2.8) based on a curated list of cell type-specific genes representing a combination of previously published genes with cluster-associated genes (Fig. S2, S8). For cells that were classified as “unknown” they were assigned the cell type that was most prominent in the corresponding cell cluster. Differential gene expression was performed through Seurat using MAST (https://genomebiology.biomedcentral.com/articles/10.1186/s13059-015-0844-5). Genes were determined to be significant if their adjusted (FDR) p-value was below 0.05. Gene set enrichment analysis in Fig. 1, Fig. 6, Fig. S4, Fig. S6) was performed using EnrichR software^77^ (maayanlab.cloud/Enrichr/) on the gene set comprising co-expression modules as identified by k-means clustering. Gene set enrichment analysis in Fig. 2 was performed using FGSEA^78^ using log fold changes as the ranking metric. Upstream regulator analysis was performed in Ingenuity Pathway Analysis software (Qiagen) on the gene set representing all differentially expressed genes (adjusted p-value <0.05) according to cell type.

To assess the biological relevance of ZIKV reads represent biologically relevant intracellular viral RNA in the iPSC-derived cell dataset, we sought to characterize ZIKV-infected cells in several ways. First, we compared gene expression profiles between infected cells and non-infected bystanders. By systematically varying the threshold for calling cells ZIKV-infected while quantifying the resulting number of DE genes, we determined a threshold of 9 UMI-distinct ZIKV reads per cell resulted in the largest number of DE genes in the differentiated condition. We used flow cytometry to confirm this estimate of the rate of ZIKV-infected cells approximated that measured by ZIKV envelope protein expression (Fig. S8B-C).

### QUANTIFICATION AND STATISTICAL ANALYSIS

For qRT-PCR gene abundance analysis, statistical tests were performed using Prism 9.0 software (GraphPad). Data are presented as mean across all measurements ± standard deviation (SD) unless otherwise stated. Statistical significance was determined using a two-tailed Student’s t test. P-values are indicated in figure legends; a p-value of <0.05 was considered statistically significant. For scRNA-seq data, statistical analysis was performed using RStudio (R Core Team; v2023.03.0) and p-values in differential expression analysis were adjusted for multiple comparisons using the Benjamini-Hoffberg method. In this study, n is determined as the number of independent biological replicates (not technical replicates) utilizing identical experimental conditions, or the number of independent experiments, as indicated in figure legends. For human primary fetal cells, a single well was considered a biological replicate when multiple wells were derived from the same initial donor (Figs. 4, S4). For iPSC-derived cells, a single well was considered a biological replicate when multiple wells were derived from the same initial differentiation (e.g. Fig. S7C).

### FLOW CYTOMETRY AND FLUORESCENCE ACTIVATED CELL SORTING

Cells were released from the dish by Accutase digestion, then triturated to produce a single cell suspension. For cell sorting, cells were resuspended in DPBS (without calcium or magnesium) containing 0.2% bovine serum albumin (BSA; Sigma-Aldrich # A7906), blocked with 2% rat serum (MilliporeSigma R9759) for 30 minutes, then incubated in viability dye (eFluor 780, ThermoFisher 65-0865-14) for 30 minutes, then incubated in antibody cocktail containing anti-CD24 (BD Pharmingen 56164) anti-CD44 (555478), and anti-CD184 (560936) for one hour at 4 degrees C. After rinsing three times in DPBS+BSA buffer, cells were sorted using a FACSAria II (BD Biosciences) following the protocol outlined in Figs. S2E and S7B.

For flow cytometry, cells were rendered into suspension as above, then fixed using 4% formaldehyde (VWR #87001-890) in PBS at room temperature for 30 minutes. Antibody staining was performed as above using anti-flavivirus envelope antibody (4G2, Absolute Antibody Ab00230) and flow cytometry was performed on a LSR II cytometer (BD Biosciences). Data analysis was performed with FlowJo software (v. 10.8).

### IMMUNOFLUORESCENCE AND IMAGE ANALYSIS

Primary human fetal tissue was fixed for one week in 4% formaldehyde (VWR #87001-890) in PBS at room temperature, then embedded in paraffin. Sections were cut at 4 μm thickness and mounted on glass slides. After deparaffinization (xylene) and rehydration (100%, 95% ethanol solution in water), antigen retrieval was performed using EDTA at 99°C for 20 minutes. Nonspecific antibody binding was blocked using blocking solution containing 2.5% normal donkey serum (Sigma-Aldrich # D9663), 2.5% normal goat serum (Jackson ImmunoResearch #005-000-121), 0.2% bovine serum albumin (BSA; Sigma-Aldrich # A7906), and 0.1% triton X-100 (Thermo Fisher #AC327371000) in phosphate-buffered saline (PBS).

Cells from tissue culture were grown on glass coverslips as above, rinsed 1x with DPBS and fixed in 4% paraformaldehyde at room temperature for 30 minutes. Fixed cells were rinsed in PBS and placed in blocking solution at room temperature for one hour, followed by incubation in primary antibody at 4°C overnight. The following day, cells were rinsed 3x with PBS and then incubated in isotype-specific, fluorophore-coupled secondary antibody at room temperature for two hours in the dark. Cells were then incubated in 200 nM DAPI (Thermo Fisher #D1306) for five minutes at room temperature, rinsed 3x with PBS, and mounted on glass slides in ProLong Gold mounting medium (Thermo Fisher #P36934). Images were acquired on a Nikon Eclipse Ti confocal microscope via a 20x air or 60x oil immersion objective using Nikon confocal software (NIS Elements).

Image analysis was performed in CellProfiler software (www.cellprofiler.org)^79^. In CellProfiler, automated image segmentation was used to establish contours representing the nucleus and soma of individual cells in order to calculate the nuclear-to-cytosolic ratio of intensity for IRF3 and NF-κB. To determine cell identity and ZIKV positivity, automated measurements of intensity were made for nuclear Sox-2 (NSCs) or olig-2 (oligodendrocyte lineage), cytosolic beta3-tubulin (neurons) or GFAP (astrocytes), or cytosolic 4G2 (ZIKV). Population-based manual thresholding was performed, guided by visual inspection of thresholding results, to identify cell types and ZIKV-positive and -negative cells.

### WESTERN BLOT ANALYSIS

Cells were lysed in modified radioimmunoprecipitation (RIPA) buffer, consisting of 150 mM NaCl, 50 mM Tris-HCl (pH 7.6), 1% triton x-100, 0.5 sodium deoxycholate) with freshly added Halt protease and phosphatase inhibitor cocktail (Thermo Fisher #78446) and okadaic acid (Calbiochem #49-560-4100). Protein concentrations were measured using the Pierce BCA protein assay kit (Thermo Fisher #23225) and 10 μg protein was subjected to SDS-PAGE and blotted onto Immobilon-P PVDF membranes (Millipore #ISEQ00010). Nonspecific antibody binding was blocked with incubation in a solution of 2% BSA (Sigma-Aldrich # A7906), 2% normal donkey serum (Jackson ImmunoResearch #005-000-121) in tris-buffered saline. Antibody labeling was assessed using SuperSignal West Pico chemiluminescence assay (ThermoFisher #34578) with image capture and analysis by ChemiDoc MP imager with ImageLab software (BioRad).

### INNATE IMMUNE STIMULATION

To achieve RIG-I stimulation, cells were transfected with immunostimulatory RNA consisting of poly-U/UC 5’ppp RNA^46,80^ using TransIT-mRNA kit (Mirus Bio #MIR 2225) following the manufacturer’s instructions. As a control, nonstimulatory RNA^80^ was transfected using the same method. To achieve TLR3 stimulation, poly(I:C) RNA was added directly to medium, and compared to no treatment (control).

### DUAL REPORTER LUCIFERASE ASSAY

Cells were co-transfected with a plasmid harboring the IFN-β-Firefly luciferase construct (IFN-Beta_pGL3)^81^ and another containing a Renilla luciferase construct (pRL) at a ratio of 40:1, (4ng IFN-β-Firefly luciferase DNA per million cells) using a TansIT-LT1 kit (Mirus Bio #MIR 2304). After 24 hours of recovery, cells were stimulated and eight hours after innate immune stimulation, cells were harvested and lysates were prepared using a Dual-Luciferase reporter assay system (Promega #E1960) with passive lysis buffer, according the manufacturer’s protocol. Luciferase activity was imaged using an automated plate reader.

### QUANTITATIVE PCR

Cells were lysed in RLT buffer (Qiagen # 79216) and total RNA was extracted from cells using the RNeasy Mini kit (Qiagen # 75144) according to the manufacturer’s instructions. cDNA was synthesized with iScript Select cDNA Synthesis Kit (Bio-Rad 1708890). cDNA was synthesized using a combination of random primer mix and oligo dT provided. qPCR was performed with 1:40 dilution of cDNA, SYBR Green Master Mix (AppliedBiosystems), and forward/reverse primers at 5µM and run on an AppliedBiosystems QuantStudio5. Relative gene expression was calculated by normalizing gene expression to reference genes *RPL13a* and *UBC*, to facilitate direct comparison between undifferentiated and differentiated cultures and across cell types^82^. Primer sequences are listed in Supplemental Table 12.

## Acknowledgments

We are grateful to the members of the flaviclub including Josh Ames, Julie Eggenberger, Emmanuelle Genoyer, Andrew Gustin, Nika Hajari, Tony Muruato, Brittany Ulloa, Alexis,. We acknowledge the hard work and careful dissection of fetal tissue by the University of Washington Birth Defects Research Laboratory (BDRL), which includes Kimberly Aldinger, Dan Doherty, Ian Phelps, Jennifer Dempsey, Yasmeen Otaibi, and Lucinda A. Cort.

Funding was provided by NIH under NIAID Grant #K08AI150996 (C.S.), NICHD R24HD000836 (I.A.G.), NINDS R01 AG080585, K01 AG059841 (J.E.Y.), and XXX (M.G.). Additional funding was provided by the University of Washington Center for Innate Immunity and Immune Disease and the Washington National Primate Center.

