## Supplemental Figures for "The Human Neural Cell Atlas of Zika Infection in developing human brain tissue: viral pathogenesis, innate immunity, and lineage reprogramming"

**Supplemental figure index:**


| Fig | Title | Pg |
|---|---|---|
| S1 | ZIKV has broad tropism in human primary fetal brain cell types | 4 |
| S2 | Classification and relative abundance of cell types in human primary brain explants | 5 |
| S3 | Detection of ZIKV RNA in human fetal brain explant scRNAseq data | 6 |
| S4 | Type I interferon signaling dominates ZIKV-induced gene expression changes in human fetal tissue infected by ZIKV virus | 7 |
| S5 | Astrocytes engage NF-κB signaling to produce interferon beta in response to ZIKV infection | 8 |
| S6 | Universal ZIKV-driven transcriptional responses in human primary fetal explants reflect apoptosis, cellular stress, and type I interferon signaling | 9 |
| S7 | Differentiation of human IPSC-derived NSCs into neurons and astrocytes, expressing mature cell markers. | 10 |
| S8 | Genetic markers used to classify cell types and reads mapping to the ZIKV genome in iPSC-derived neural cell cultures by single cell RNA sequencing | 11 |
| S9 | Sensitivity analysis comparing RNA sequencing and flow cytometry readouts of ZIKV positivity in IPSC-derived neural cells | 12 |
| S10 | IFN-β dominates the transcriptional response to ZIKV infection in iPSC-derived neural cells | 13 |
| S11 | Gene expression changes reflecting apoptosis signaling are engaged with increasing levels of ZIKV copy number across cell types. | 14 |

**Supplemental table index:**


| Tb | Title | Pg |
| --- | --- | --- |
| ST1 | Human primary fetal brain tissue gestational age and sex | 3 |
| ST2 | Differential gene expression by cell type in response to ZIKV-Brazil, ZIKV-Cambodia, and IFN-β in human fetal brain explants. Data from heatmap in Fig. 1E, UpSet plot in Fig. S4A | -- |
| ST3 | Over-representation analysis of differentially expressed genes in ZIKV Brazil or ZIKV Cambodia infection across cell types in human fetal brain explants. Related to Fig. 1F, Fig. S4B | -- |
| ST4 | Differential gene expression according to cell type in ZIKV+ versus ZIKV- cells in human fetal brain explants. Data from heatmap in Fig. 3A | -- |
| ST5 | Gene set enrichment in ZIKV+ versus ZIKV- cells according to cell type in human fetal brain explants. Related to Fig. 3A | -- |
| ST6 | Differential gene expression in ZIKV-positive and ZIKV-negative cells versus MOCK condition according to cell type in human fetal brain explants. Data from heatmap in Fig. S6 | -- |
| ST7 | Over-representation analysis of differentially-expressed genes in ZIKV-positive and ZIKV-negative cells versus MOCK in human fetal brain explants. Related to Fig. S6 | -- |
| ST8 | Differential gene expression according to cell type in response to ZIKV-Brazil and IFN-β in IPSC-derived neural cells. Data from heatmap in Fig. 4G, UpSet plot in Fig. S9A. | -- |
| ST9 | Differential gene expression in ZIKV+ and ZIKV- cells versus MOCK condition according to cell type in IPSC-derived neural cells. Data from heatmap in Fig. 5A. | -- |
| ST10 | Over-representation analysis of differentially-expressed genes in ZIKV-positive and ZIKV-negative cells versus MOCK in IPSC-derived neural cells. Related to Fig. 5 | -- |
| ST11 | Over-representation analysis of differentially-expressed genes in ZIKV-positive neurons versus MOCK in IPSC-derived neural cells. | -- |
| ST12 | Primer sequences used in RT-PCR assays | |
| ST13 | Antibodies used for immunochemical assays | |

**Table ST1. Human primary fetal brain tissue gestational age and sex**

| Donor Identity | Gestational age | Sex | Experiments in which tissue was used |
|---|---|---|---|
| Donor 1 | 117 days | M | 1. Single cell RNA sequencing<br>2. IRF3 and NFκB nuclear translocation (Fig. 2) |
| Donor 2 | 108 days | M | 1. Zika tropism (Fig. S1)<br>2. IRF3 and NFκB nuclear translocation |
| Donor 3 | 89 days | F | 1. Zika tropism (Fig. S1) |

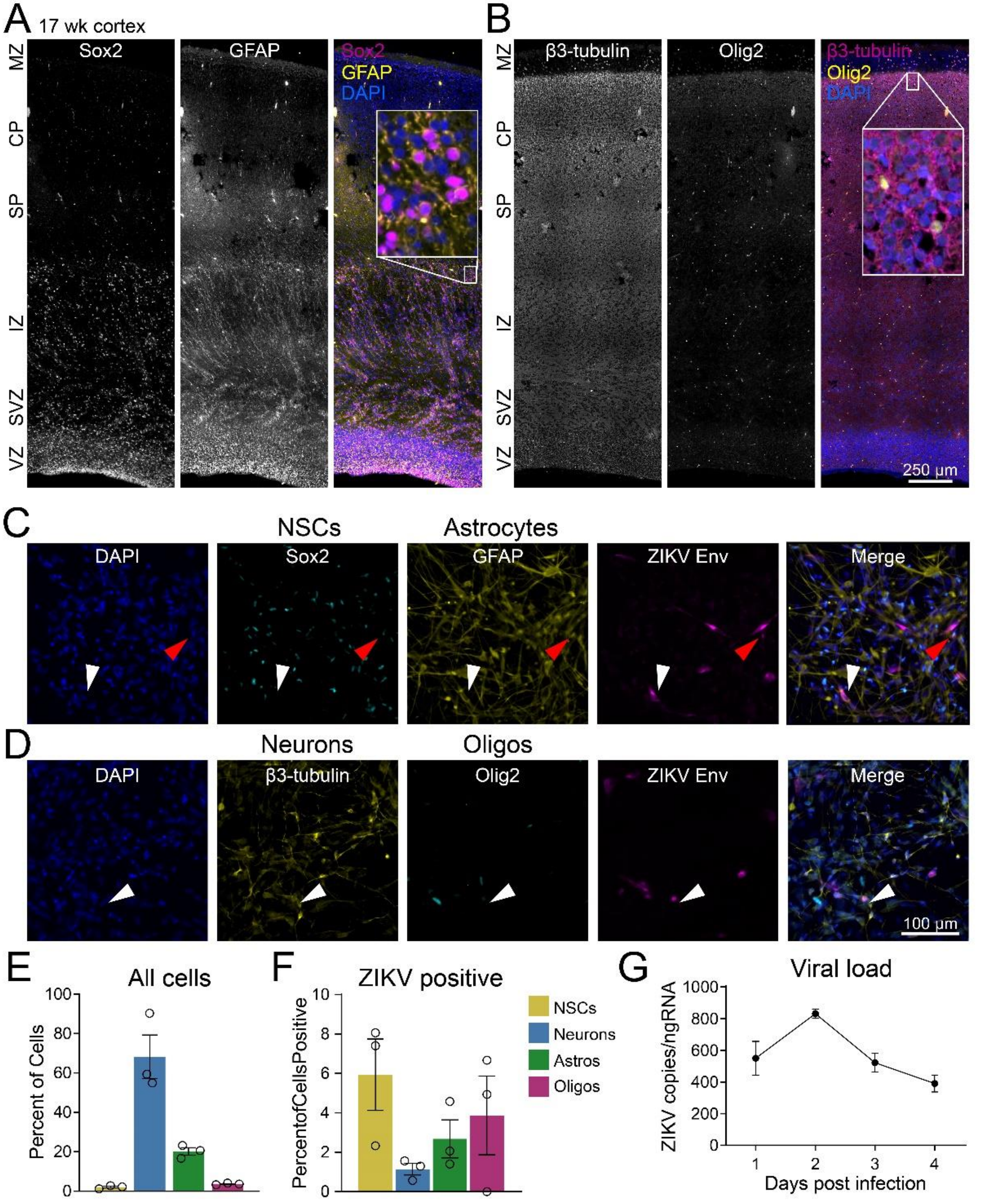


**Figure S1. ZIKV has broad tropism in human fetal brain cell types.** (A) Full-thickness section of developing fetal cortex from a 117-day male conceptus (donor 1) with immunolabeling of neural progenitors (Sox-2, yellow) and astrocytes (GFAP, magenta). Note that radial glia also express GFAP. (B) Adjacent slice from the same brain as A, with immunolabeling for neurons (β3-tubulin, magenta) and oligodendrocyte lineage cells (Olig2, yellow). (C) Two-dimensional mixed primary cultures (donor

1) demonstrating Zika infection (Env, magenta) of neural progenitors (Sox2, cyan) and astrocytes (GFAP, yellow). MOI=20, 2 days post-infection. Red arrowhead, ZIKV-positive neural progenitor; white arrowhead, ZIKV-positive astrocyte. (D) Immunostaining of cultures from the same experimental conditions demonstrating Zika infection of neurons (β3-tubulin, yellow) and oligodendrocyte lineage cells (olig2, magenta). White arrowhead, ZIKV-positive neuron. (E) Quantification of cell types as a percentage of total cells in the cultures shown in C and D. (F) Quantification of the percentage of each cell type infected by Zika virus, as measured by positive Env staining. (G) RT-PCR using ZIKV-specific primers demonstrates evolving viral load during a four-day time course of infection (MOI=20).

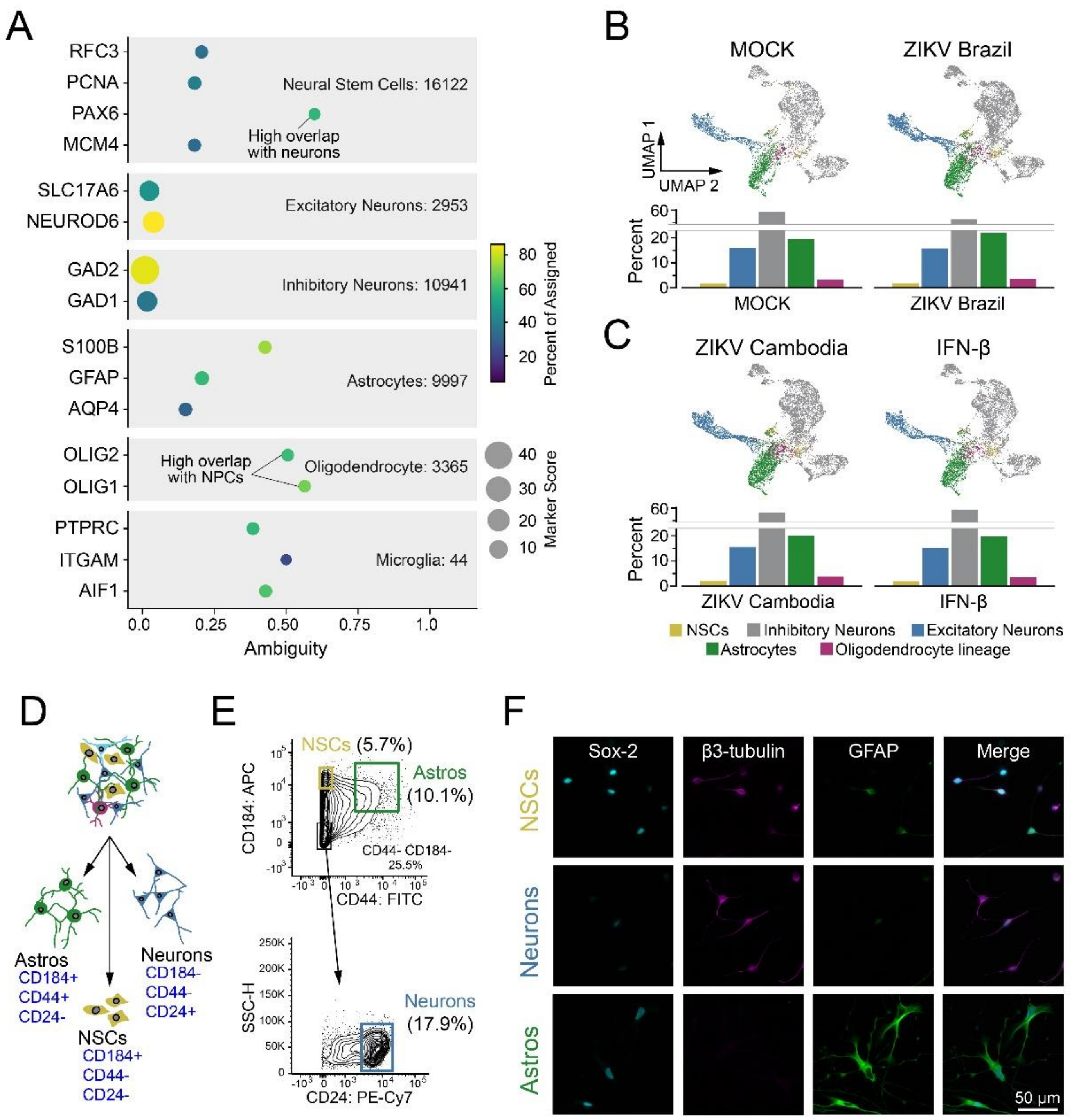


**Figure S2. Classification and relative abundance of cell types in human primary brain explants.** (A) Cell classifiers representing the combination of: (1) a list of curated genes based on published markers (e.g. olig2) and (2) automated identification of marker genes using Garnett software (e.g. SLC17A6). (B) Uniform manifold approximation and Projection (UMAP) plots (top row) indicate clustering of neural progenitors (gold), excitatory neurons (blue), inhibitory neurons (grey), astrocytes (green), and oligodendrocyte lineage cells (purple). Bottom row, relative abundance of identified cell types in the MOCK and ZIKV-Brazil conditions. (C) UMAP (top) and relative abundance (bottom) of cell types in ZIKV-Cambodia and IFN-β conditions. (D-E) Fluorescence activated cell sorting (FACS) approach (D) and results (E), based on protocols established in induced neural progenitor cells (Yuan et al.), confirms rough proportions of cell types identified by cell clustering in scRNAseq data. Compare to Fig. S7B (cell sorting in iPSC-derived neural cells). (F) Immunofluorescence of purified cell populations demonstrating enrichment of Sox2 (cyan) in NSCs (top), β3-tubulin (magenta) in neurons (middle), and GFAP (yellow) in astrocytes.

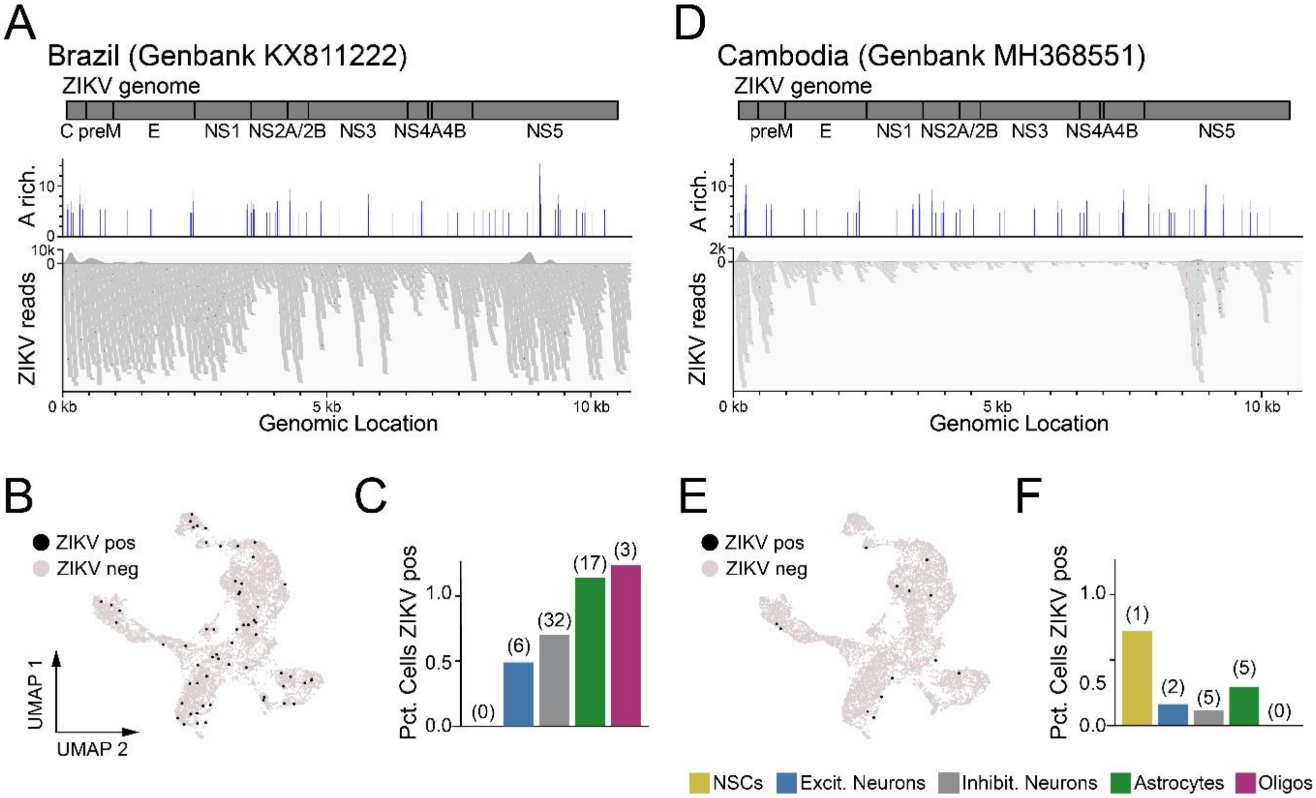


**Figure S3. Detection of ZIKV RNA in human fetal brain explant scRNAseq data using scPathoQuant.** (A) Top, architecture of the ZIKV genome. Middle, bar plots showing adenosine-rich segments of the Brazilian strain of ZIKV (GenBank no. KX811222). Enrichment was calculated as significant increase in adenosine over a 30-base segment, as compared to the whole genome. Bar height represents the percentage of adenosine bases in a 30-base segment and is only plotted for segments where significant enrichment was found. Bottom, distribution of reads (mapped by Interactive Genome Viewer) identified in all single cell RNA sequencing samples corresponding to Brazilian strain. (B) Uniform manifold approximation and projection (UMAP) plot of cells containing ZIKV reads (black points) and those without (grey points) in ZIKV-Brazil data set. (C) Frequency of ZIKV-positive cells (percent positive) according to cell type in ZIKV-Brazil dataset. Numbers above bars indicate total number of cells ZIKV positive for the indicated cell type. (D) Architecture of the ZIKV genome (top), A-enrichment map (middle), and reads mapping to the Cambodian strain of ZIKV (GenBank no. MH368551; bottom), as in A. (E-F) UMAP (E) and percentages (F) of cells containing ZIKV reads in the ZIKV-Cam dataset, as in B-C.

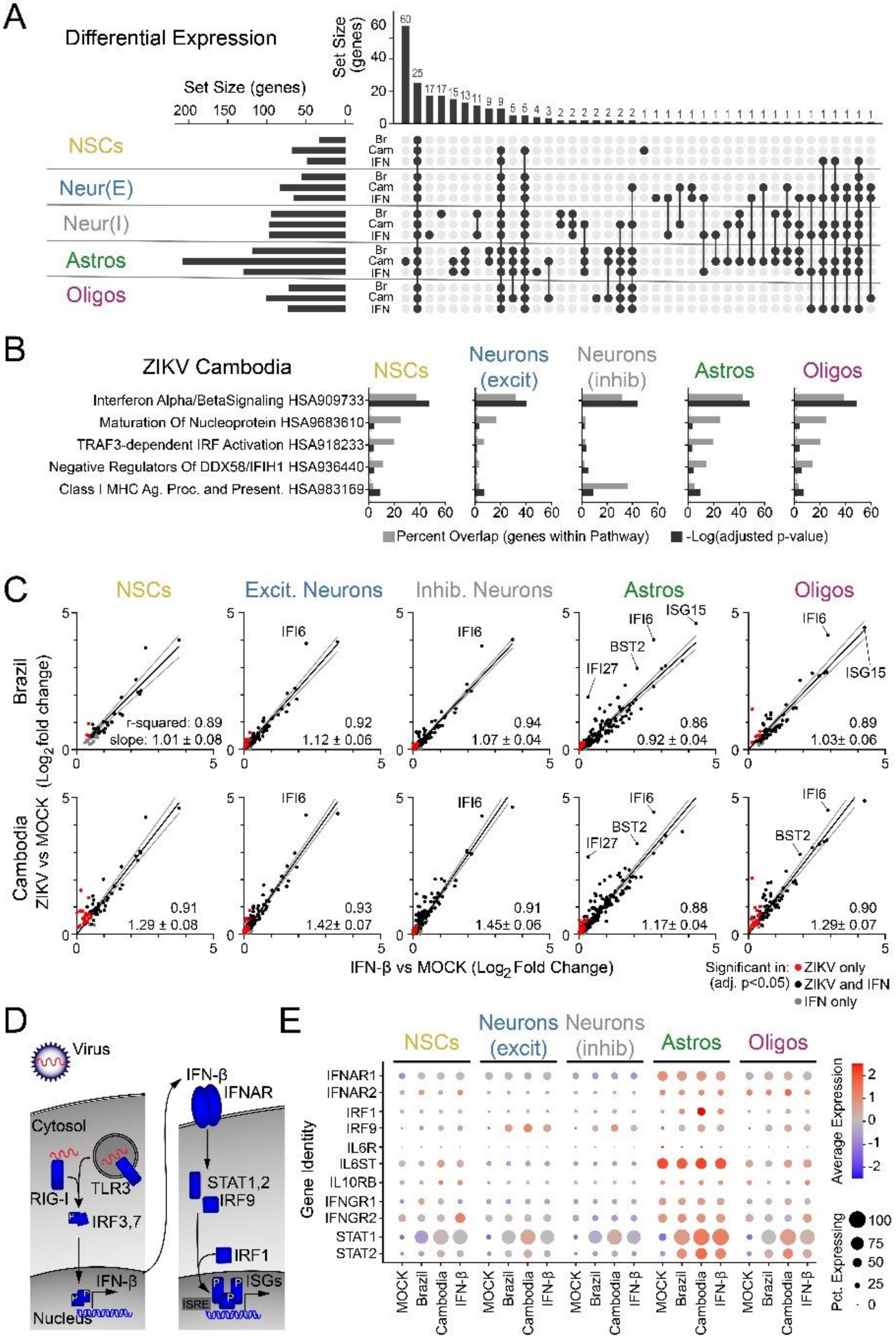


**Figure S4. Type I interferon signaling dominates ZIKV-induced gene expression changes in human fetal tissue infected by ZIKV virus.** Related to Fig. 1-2. (A) UpSet plot indicating overlap in gene expression changes (adj. p-value <0.05, absolute log fold change >1.2) induced across cell types by ZIKV infection (BR=Brazil, CAM=Cambodia) or IFN-β treatment (100 IU for 6 hours). (B) Gene ontogeny (GO) signatures identified by Gene Set Enrichment Analysis (GSEA) for each cell type based on significant DE genes in response to ZIKV Cambodia. (C) Scatterplots (points=genes) representing $\log_2$ fold change in ZIKV infection (y-axis) versus the $\log_2$ fold change in response to IFN-β treatment (x-axis). Top, ZIKV-Brazil; bottom, ZIKV-Cambodia. Genes were included if they were significantly upregulated in ZIKV treatment only (red), IFN-β treatment (grey), or both conditions (black). Most genes fall close to a linear relationship (black line=simple linear regression constrained through the origin; slope (+/- 95% confidence interval) is shown in the lower right-hand corner of each graph) reflecting the similarity of ZIKV-induced gene expression to IFN-β-induced gene expression. R-squared value was calculated from linear regression of the same points without constraining the line to the origin. (D) Schematic of canonical type I interferon signaling indicating the relationship between the four TFs identified in Fig. 2A. (E) Dot plot representation of treatment-specific gene expression for type I interferon receptors (IFNAR1 and IFNAR2) as well as additional innate immune signaling pathways of interest. Color scale represents normalized expression level across all genes and all conditions represented in the plot.

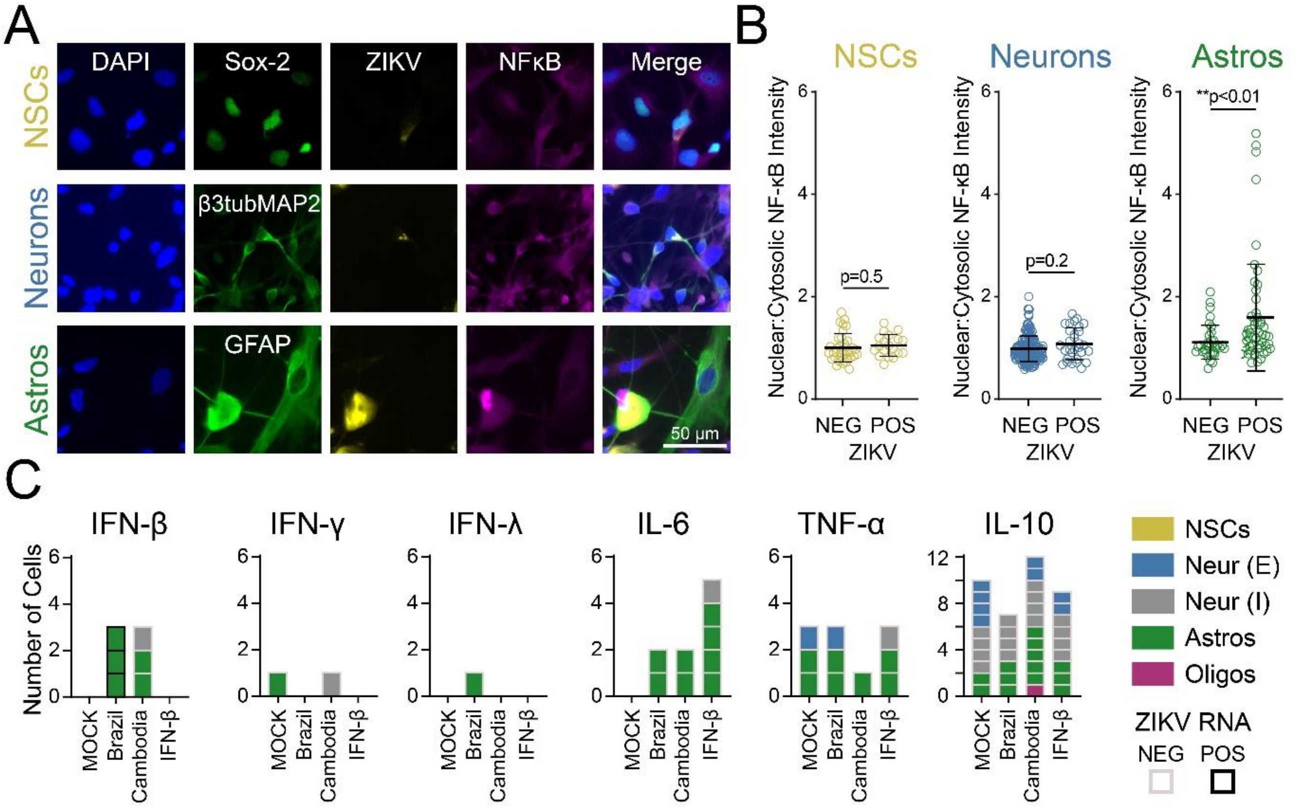


**Figure S5. Astrocytes engage NF-κB signaling to produce interferon beta in response to ZIKV infection.** Related to Fig. 2. (A) Immunocytochemistry (ICC) of innate immune activation, indicated by nuclear translocation of NF-κB, in ZIKV-infected mixed human fetal primary brain cells (Donor 2). (B) Quantification of NF-κB nuclear translocation, measured as the ratio of nuclear-to-cytosolic immunofluorescence, demonstrated innate immune activation only in a subset of ZIKV-positive astrocytes. N=1 experiment (3 coverslips quantified per cell type) from donor 2, representative of two similar experiments from two donors. Error bars reflect mean ± SD; p-value calculated by Kolmogorov-Smirnov test. (C) Cytokine expression in human primary fetal brain cells represented as the number of cells containing reads for the identified cytokine, according to treatment condition (x-axis), cell type (color) and ZIKV infection (black or grey outline). The following type I interferons did not have any detectable reads in any cells: IFN-α1, α2, α4, α5, α8, α10, α13, α14, α17, α21.

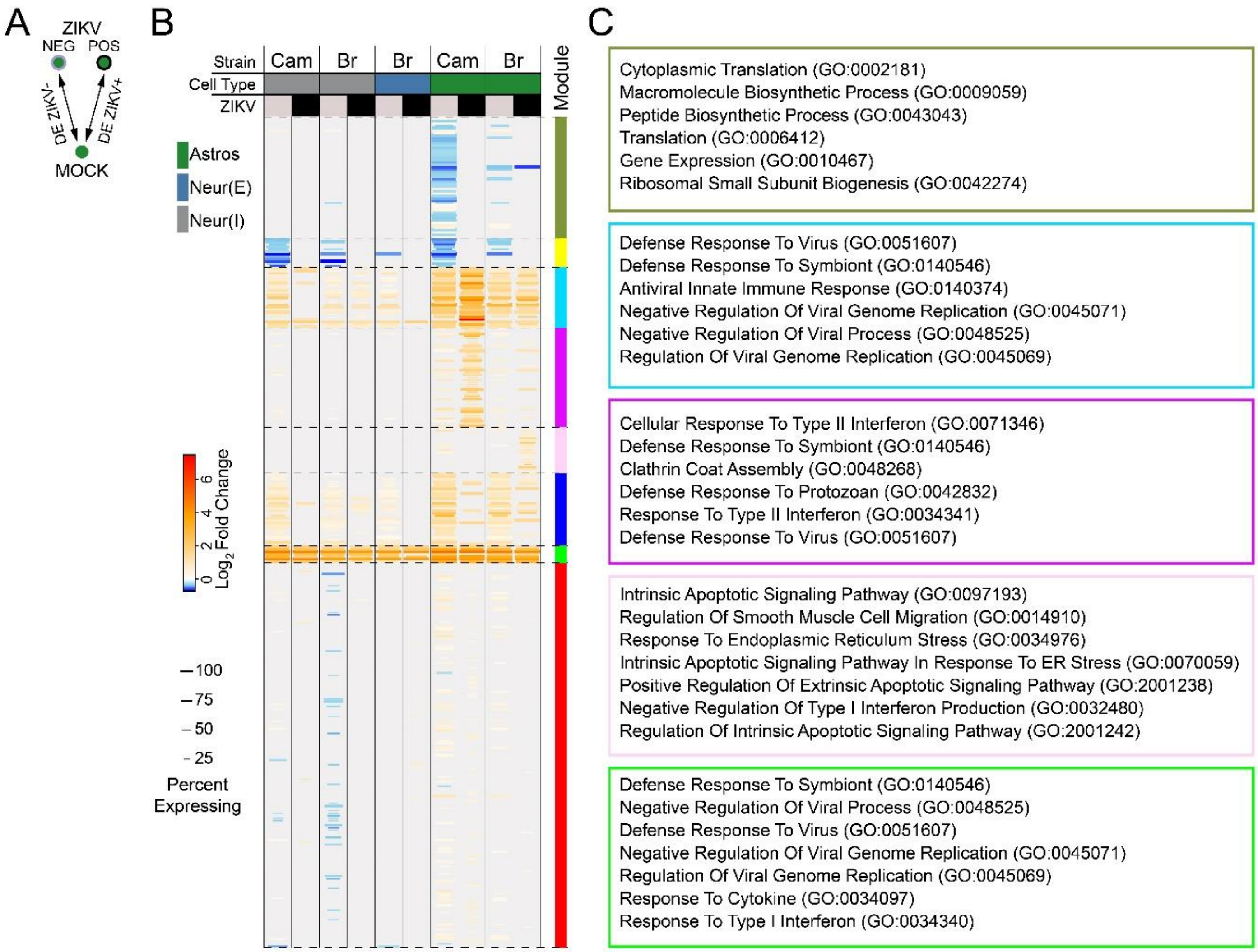


**Figure S6. Universal ZIKV-driven transcriptional responses in human primary fetal explants reflect apoptosis, cellular stress, and type I interferon signaling.** (A) Schematic of DE comparison. (B) Heatmap of DE genes (rows=genes) grouped by viral strain, cell type (color column headings) and ZIKV-positivity (black or grey column headings). Clustering of genes by K-means identified modules of DE genes (right color bar) representing several categories of response. Some gene modules are universal across cell types (e.g., yellow, red modules), some modules are specific to differentiated cells (light blue module), and some modules are specific to neurons (magenta and dark green modules). Only significant (adj. p-val<0.05) DE genes are shown. (**C**) Overrepresentation analysis of DE gene modules (identified by color box) demonstrating functional pathways enriched within each module.

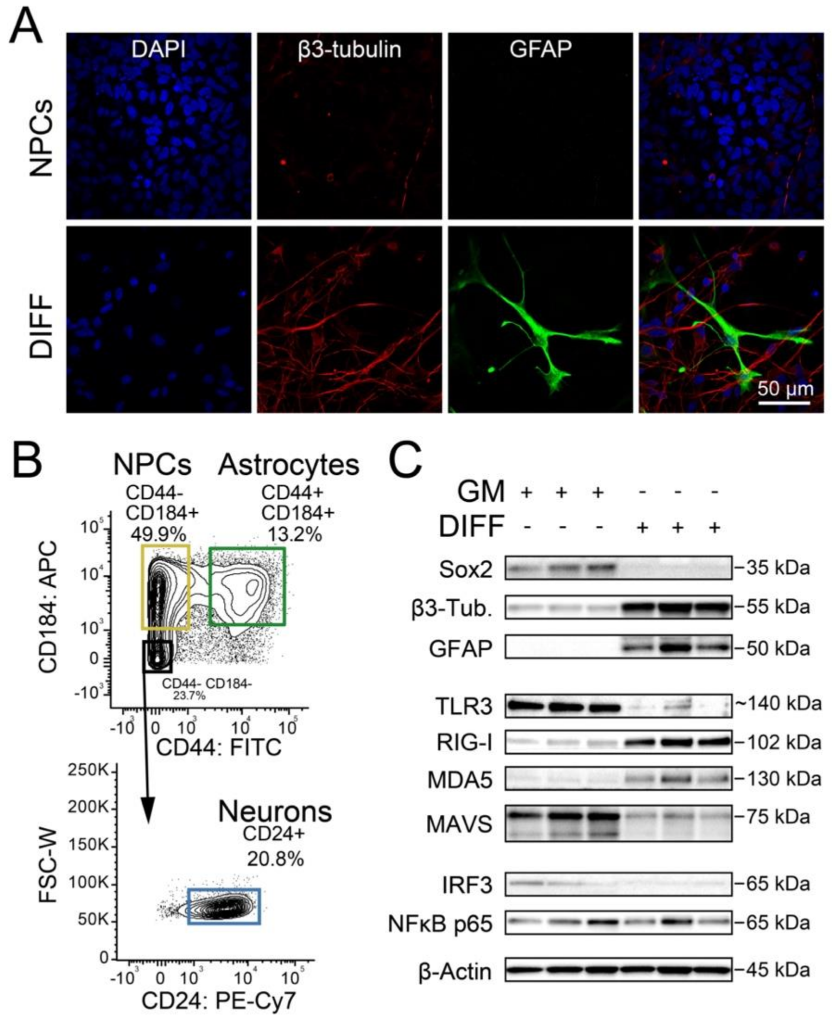


**Figure S7. Differentiation of human IPSC-derived NSCs into neurons and astrocytes, expressing mature cell markers.** (A) Neural progenitor cultures ("STEM", top) derived from IPScs can be stably passaged when maintained in growth medium containing basic fibroblast growth factor (bFGF). Bottom, differentiated cultures were produced by removal of bFGF, followed by a minimum of six weeks of growth in the presence of neurotrophic growth factors (brain-derived neurotrophic factor, BDNF and leukemia inhibitory factor, LIF). Differentiated cultures contained mature neurons (β3-tubulin, red) and astrocytes (GFAP, green). (B) Utilizing an established protocol for fluorescence activated cell sorting, purified populations of NSCs, neurons, and astrocytes can be isolated. (C) Western Blot demonstrating resting expression levels of innate immune signaling components for detecting RNA viruses, in triplicate cultures of progenitors (three left lanes) and differentiated cells (three right lanes).

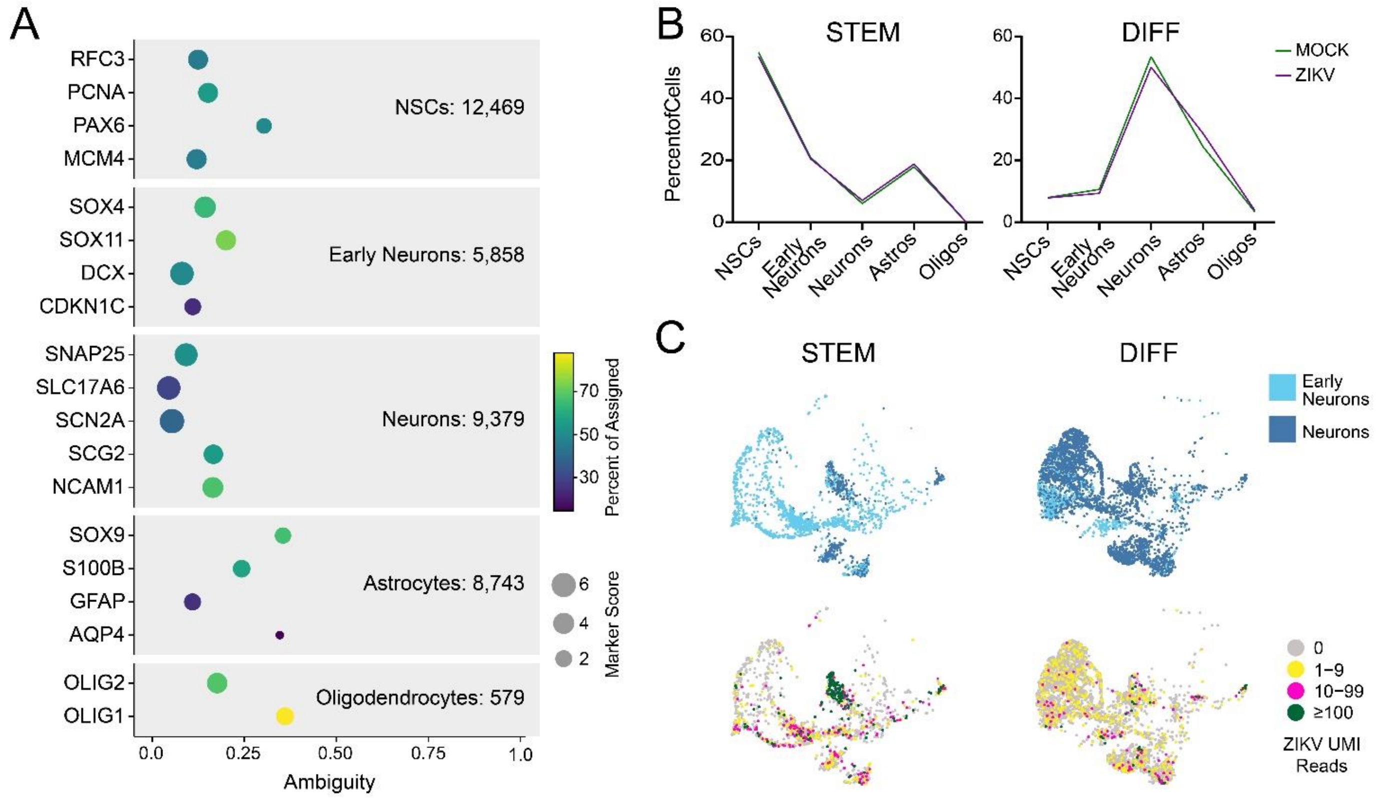


**Figure S8. Genetic markers used to classify cell types and reads mapping to the ZIKV genome in iPSC-derived neural cell cultures by single cell RNA sequencing.** Cell classifiers built using Garnett software, using a combined list of curated genes based on published markers (e.g. olig2) and automated identification of marker genes (e.g. SLC17A6). (B) Relative percentages of identified cell types in undifferentiated ("STEM") and differentiated cultures ("DIFF"), in MOCK and ZIKV-infected conditions. (C) Uniform Manifold Approximation and Projection (UMAP) plots demonstrating early neurons and neurons identified in STEM and DIFF conditions (top) and raw ZIKV UMI counts in the same cell types (bottom).

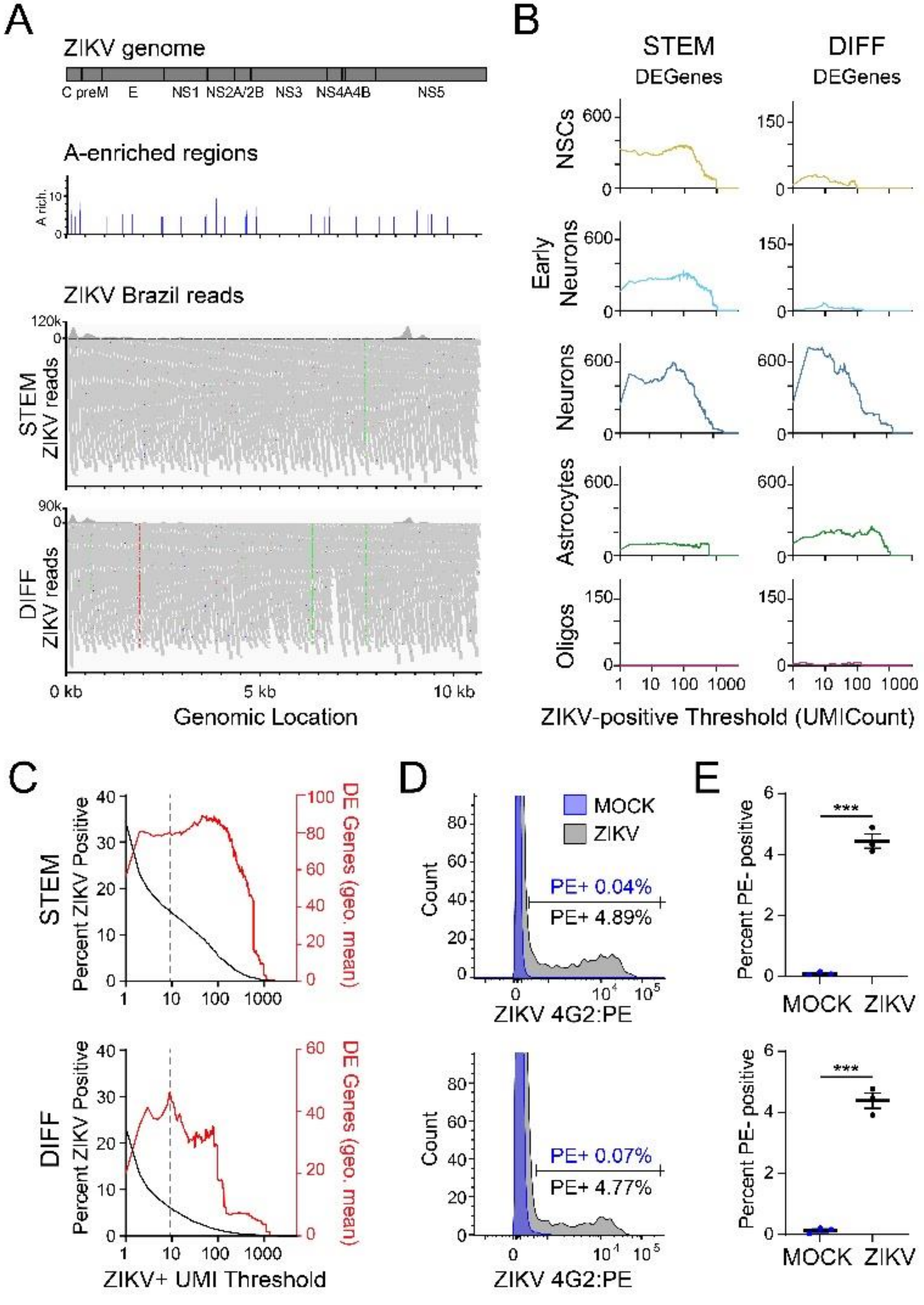


**Figure S9. Sensitivity analysis comparing RNA sequencing and flow cytometry readouts of ZIKV positivity in IPSC-derived neural cells.** (A) Top, architecture of the ZIKV genome. Middle, bar plots showing adenosine-rich segments of the Brazilian strain of ZIKV (GenBank no. KX811222). Bottom, distribution of reads (mapped by Interactive Genome Viewer) identified in all single cell RNA sequencing samples corresponding to Brazilian strain. (B-C) Using scRNAseq data, the ZIKV-positive threshold (x-axis) was defined as the number of unique molecular identifier (UMI)-tagged sequences mapping exclusively to the ZIKV genome required to call a cell "ZIKV-positive". (B) By performing DE analysis while progressively adjusting the threshold of ZIKV reads at which cells were considered positive (from 1 to 4742, the highest number of ZIKV reads in a single cell) we generated a curve representing the number of significantly (adj. p-value <0.05) DE genes between ZIKV-positive and ZIKV-negative cells at each threshold (Fig. S8C). Sensitivity analysis quantifying the number of significantly DE genes when comparing ZIKV-positive to ZIKV-negative cells at each threshold for identified cell types in progenitor (top) and differentiated cell cultures (bottom). (C) Combined sensitivity analysis analyzing the percent of all cells that are ZIKV positive (black curve, left axis) and the number of DE genes for all cell types (red curve, right axis; calculated as the geometric mean of DE genes across cell types depicted in A) for each ZIKV-positive threshold (x-axis). The largest number of DE genes in differentiated cells was observed at a ZIKV-positive UMI threshold of 9. (C-D) Estimates of ZIKV-positive cells made by flow cytometry of bulk cultures of ZIKV infection in progenitors and differentiated cultures. (C) Fluorescence intensity histogram indicating distribution of cellular labeling for ZIKV envelope protein (x-axis) in MOCK- (blue) and ZIKV-infected (blue) conditions in progenitor (top) and differentiated (bottom) cell cultures. Infection conditions were chosen to mimic the conditions used in single cell RNA sequencing experiments: MOI = 10, 42hrs. Threshold was chosen manually by visual inspection of data. (D) Summary of flow cytometry estimates of ZIKV-positive cells (n=3 independent experiments). ***p<0.001. For simplicity, we elected to use a threshold of ≥9 UMI-tagged reads per cell for both progenitor and differentiated cultures in our initial analysis (Fig. 6A).

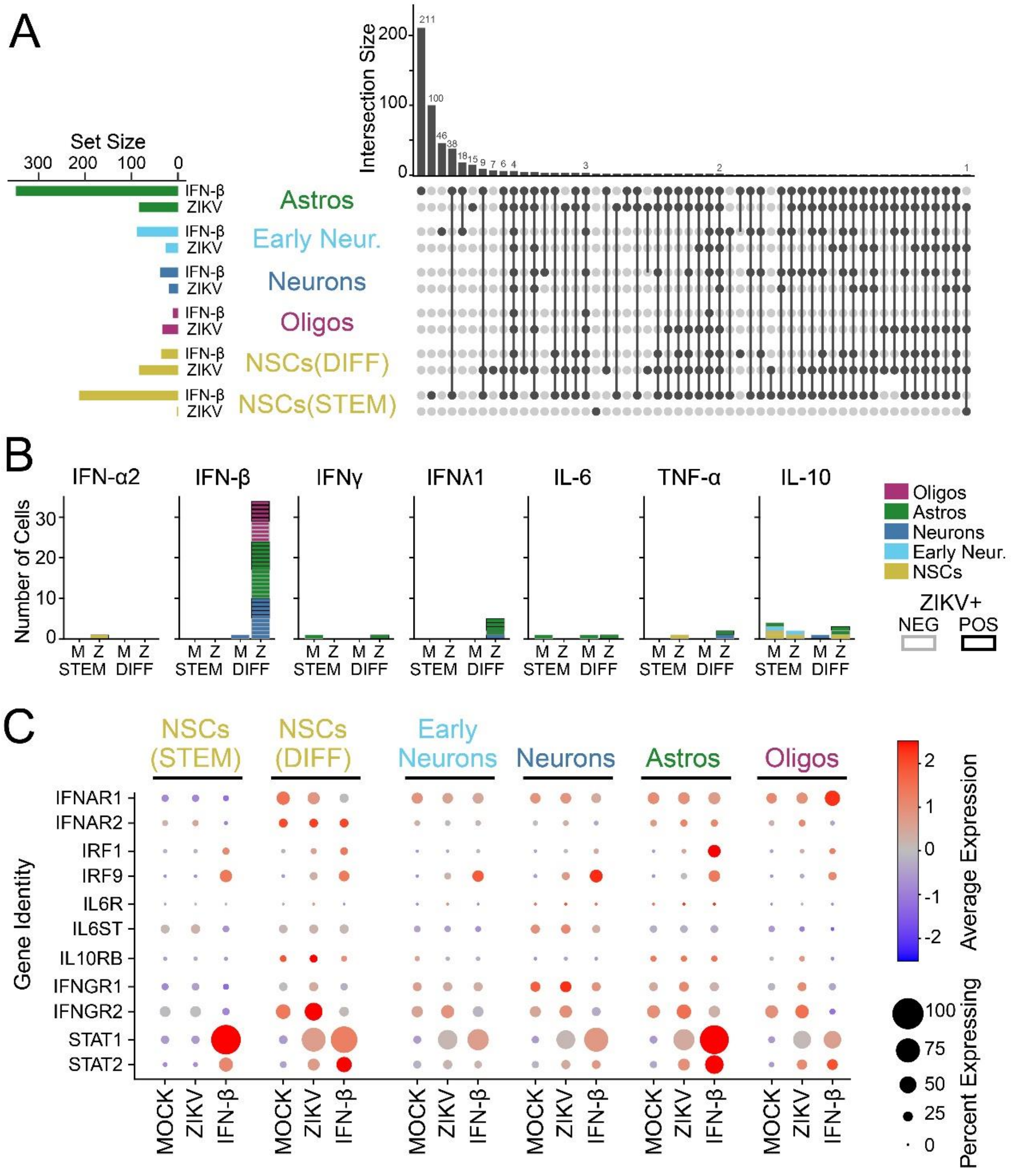


**Figure S10. IFN-β dominates the transcriptional response to ZIKV infection in iPSC-derived neural cells.** Related to Fig. 4-5. (A) UpSet plot indicating overlap in gene expression changes (adj. p-value <0.05, absolute $\log_2$ fold change >0.2) induced by ZIKV infection or IFN-β treatment (100 IU for 6 hours) across cell types and differentiation states. (B) Cytokine expression in human IPSC-derived neural cells represented as the number of cells containing reads for the identified cytokine, according to treatment condition (x-axis), cell type (color) and ZIKV infection (black or grey outline). The following type I interferons had reads detected in one or no cells across the dataset: IFN-α1, α2, α4, α5, α8, α10, α13, α14, α17, α21. (C) Dot plot representation of treatment-specific gene expression for type I interferon receptors (IFNAR1 and IFNAR2) as well as additional innate immune signaling pathways of interest. Color scale represents normalized expression level across all genes and all conditions represented in the plot.

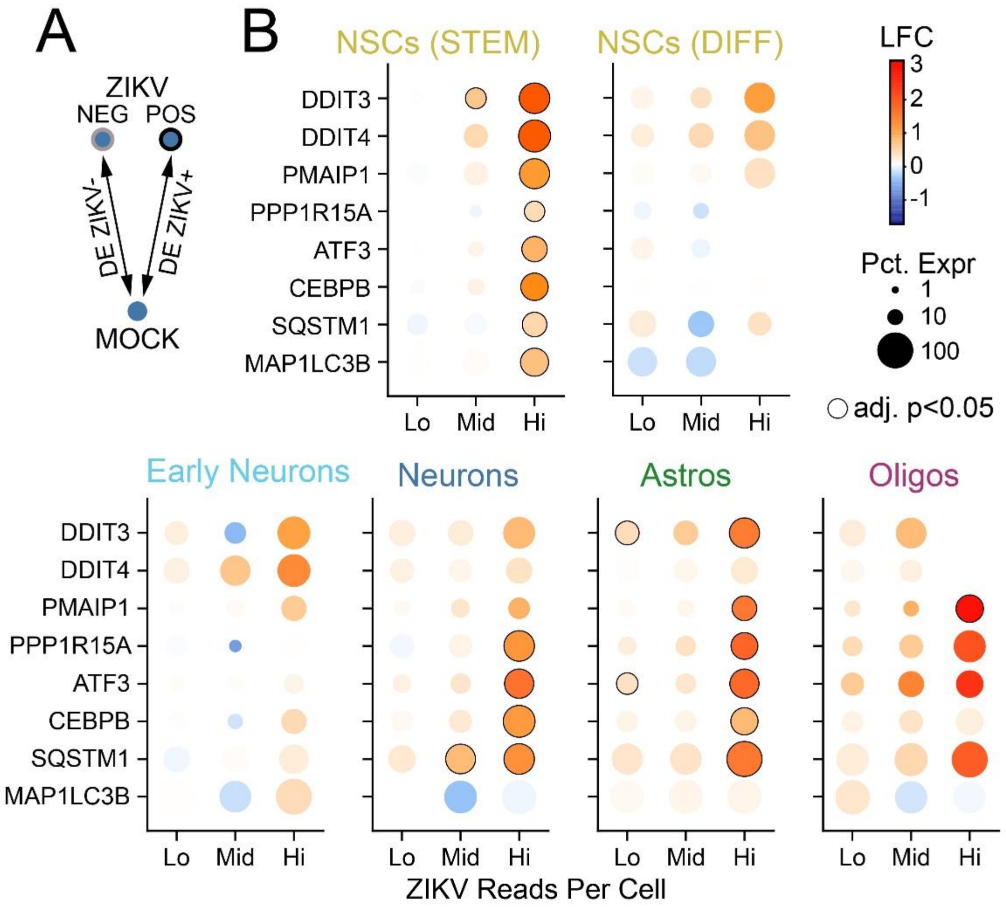


**Figure S11. Gene expression changes reflecting apoptosis signaling are engaged with increasing levels of ZIKV copy number across cell types.** (A) Schematic of DE comparison. (B) Dot plots representing gene expression changes in ZIKV-positive cells (vs MOCK) as a function of the number of ZIKV UMI reads (x-axis), for genes pertaining to apoptosis (*DDIT3, DDIT4, PMAIP1, PPP1R15A, ATF3, CEBPB*) and autophagy (*SQSTM1* and *MAP1LC3B*). Lo: 1-9 reads; Mid: 10-99 reads; Hi:>99 reads. Color: log-fold change indicated by color bar in B; size=percent of cells expressing gene.
